# Functional biogeography of marine microbial heterotrophs

**DOI:** 10.1101/2024.02.14.580411

**Authors:** Emily J. Zakem, Jesse McNichol, J.L. Weissman, Yubin Raut, Liang Xu, Elisa R. Halewood, Craig A. Carlson, Stephanie Dutkiewicz, Jed A. Fuhrman, Naomi M. Levine

**Affiliations:** Department of Global Ecology; Carnegie Institution for Science, Stanford, CA, USA; Department of Biology; St. Francis Xavier University, Antigonish, Nova Scotia, Canada; Department of Biological Sciences; University of Southern California, Los Angeles, CA, USA; Institute for Advanced Computational Science; Stony Brook University, Stony Brook, NY, USA; Department of Ecology and Evolution; Stony Brook University, Stony Brook, NY, USA; Department of Biology; The City College of New York, New York, NY, USA; Department of Earth, Atmospheric and Planetary Sciences; Massachusetts Institute of Technology, Cambridge, MA, USA; Department of Ecology, Evolution, and Marine Biology; Marine Science Institute, University of California, Santa Barbara, CA, USA; Center for Sustainability Science and Strategy; Massachusetts Institute of Technology, Cambridge, MA, USA

## Abstract

Heterotrophic bacteria and archaea (‘heteroprokaryotes’) drive global carbon cycling, but how to quantitatively organize their functional complexity remains unclear. We generated a global-scale understanding of marine heteroprokaryotic functional biogeography by synthesizing genetic sequencing data with a mechanistic marine ecosystem model. We incorporated heteroprokaryotic diversity into the trait-based model along two axes: substrate lability and growth strategy. Using genetic sequences along three ocean transects, we compiled 21 heteroprokaryotic guilds and estimated their degree of optimization for rapid growth (copiotrophy). Data and model consistency indicated that gradients in grazing and substrate lability predominantly set biogeographical patterns, and identified deep-ocean ‘slow copiotrophs’ whose ecological interactions control the surface accumulation of dissolved organic carbon.

## Introduction

The global carbon cycle plays a central role in the Earth’s climate by regulating the amount of carbon dioxide (CO_2_) in the atmosphere. Photosynthesis removes CO_2_ from the atmosphere and converts it to organic carbon, which rapidly cycles through the biosphere. Ultimately, most organic carbon is consumed by heterotrophic organisms and respired back into CO_2_. In the ocean, both photosynthesis and respiration are driven predominantly by microorganisms (*1*), and their decoupling in time and space sequesters carbon (*2*).

How effectively microorganisms can mediate ocean carbon sequestration influences Earth’s future climate trajectory. However, in the ocean biogeochemical models used for climate projections, the activities of heterotrophic Bacteria and Archaea (heterotrophic prokaryotes; hereafter, ‘heteroprokaryotes’) that consume and respire organic carbon are typically represented implicitly and simplistically (*3*). This prevents climate models from capturing potential microbial feedbacks (*4*). Simplistic resolution reflects our limited understanding of how heteroprokaryotic communities control carbon storage. Respiration in the deep ocean enhances the concentrations of dissolved inorganic carbon, but models cannot agree on the sign of the change in the organic carbon exported to the deep ocean in a warming climate (*5*). Dissolved organic carbon (DOC) is one of the largest biologically available carbon reservoirs in the earth system, and it accumulates throughout the ocean, with higher concentrations in surface waters (*6*). However, the limitations to heteroprokaryotic consumption that allow DOC to accumulate remain unclear (*7, 8*).

Before we can build a robust and dynamic heteroprokaryotic response into climate models, we must develop a mechanistic understanding of how heteroprokaryotic communities are currently structured and how this structure relates to their biogeochemical function. This is a challenging task because heteroprokaryotic communities are immensely complex, consisting of thousands of mostly uncultivated populations consuming thousands of organic substrates, many of which are uncharacterized (*9–12*). What are the broad classifications appropriate for representing heteroprokaryotic diversity quantitatively at large scales? Rapidly expanding global-scale sequencing efforts have the potential to address some of these unknowns and to provide the data needed to inform and analyze ecosystem models (*13–15*). However, new approaches are needed to fully harness these datasets and connect them with modeling frameworks (*16*).

A tractable approach for untangling heteroprokaryotic community complexity is to leverage established organizing principles. First, heteroprokaryotes have long been classified as either “copiotrophs,” with cellular machinery optimized for rapid growth when substrate is abundant, or “oligotrophs,” optimized for competitive uptake of limited substrates (i.e., high affinity) (*17–20*). This tradeoff between growth rate and substrate affinity reflects a cell’s finite proteome, and thus the amount of cellular machinery that can be allocated to a given function (*21*). This classification has been used to explain biogeographical patterns in the ocean surface: oligotrophs dominate the microbial community in subtropical ‘ocean deserts’ while copiotrophs become more prevalent in high-latitude, nutrient-rich waters (*22–24*). Second, with respect to the organic carbon substrates consumed by heteroprokaryotes, previous work has organized these substrates along an axis of “lability,” correlating with turnover rates (*6*). Due to the confounding impacts of variable energetic content, enzymatic cost, enzymatic rates, the metabolic capabilities of the local microbial community, and other factors (*7*), compounds range from highly labile, with turnover rates of minutes, to highly recalcitrant, with turnover rates of decades to millennia (*6, 25*).

We developed a global ocean ecosystem model that used these two established organizing principles to resolve heteroprokaryotic functional diversity (Fig. 1). We compared the global heteroprokaryotic biogeographical distributions predicted by the model to the coarse-grained patterns revealed from sequencing datasets from three transects that span the global ocean (Fig. 2A). Our integrated analysis of model and data identified and explained structural and functional patterns of the heteroprokaryotic metacommunity. Both model and data identified a vertical transition from predominantly oligotrophic populations in low-latitude surface waters to deep populations that grow more slowly than surface populations, but are demonstrably more copiotrophic.

**Fig. 1.**
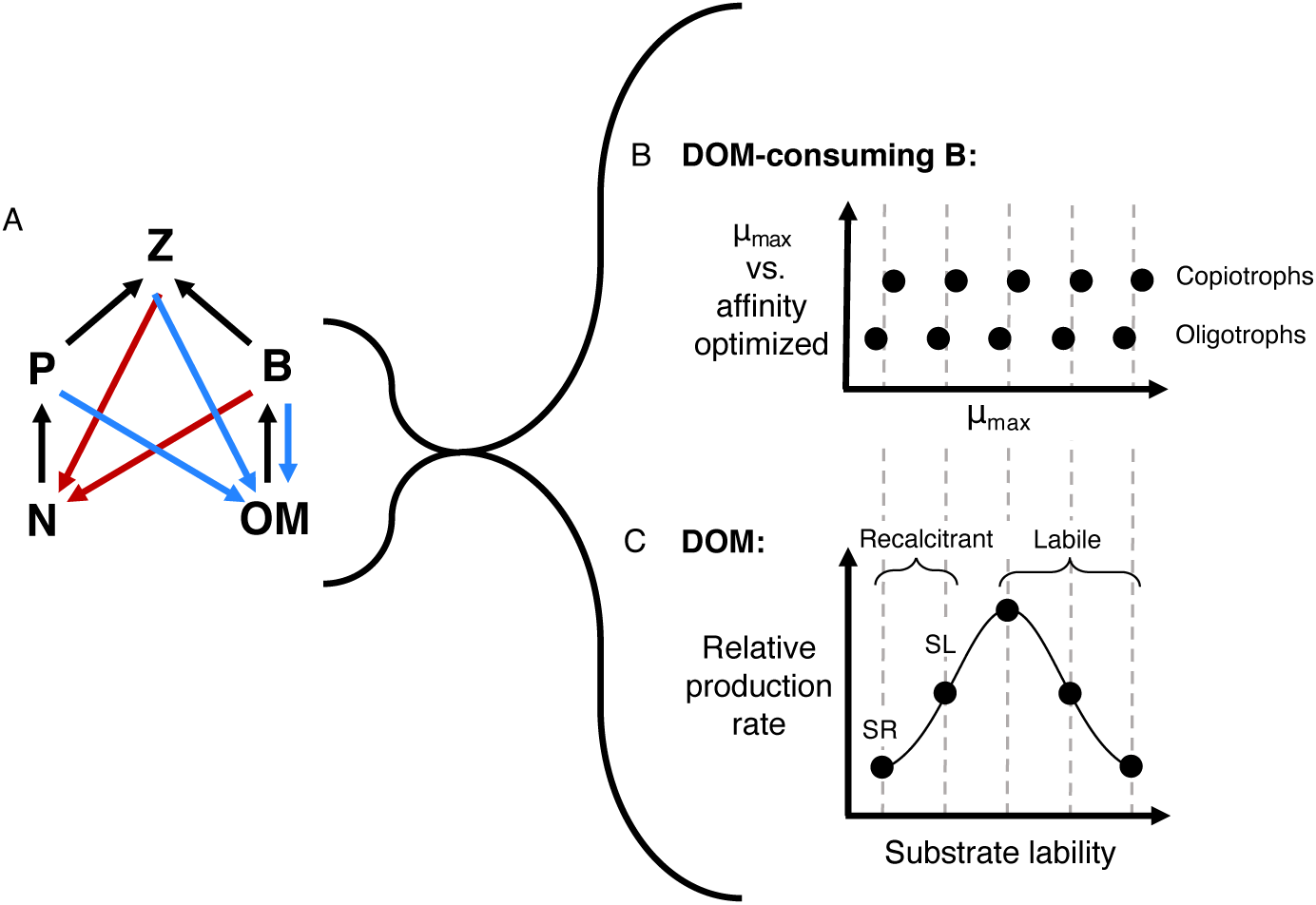
Schematic of the marine ecosystem model with heterotrophic prokaryotes (‘heteroprokaryotes’). (**A**) The model resolves populations of heteroprokaryotes (*B*, for bacteria), organic matter (*OM*), phytoplankton (*P*), inorganic nutrients (*N*), and zooplankton grazers (*Z*). Populations and substrates are connected via consumption (black arrows), respiration (red arrows), and mortality (blue arrows). (**B**) DOM-consuming heteroprokaryotic diversity is resolved with two axes: lability (x-axis), defined by setting the baseline maximum growth rate (*μ_max_*) of consumers, and a metabolic trade-off (y-axis) between maximum growth rate (copiotrophy) and substrate affinity (oligotrophy). (**C**) Total DOM production is partitioned into five classes following a lognormal distribution (*7, 33*). These five classes align with empirical and dynamical classifications (*6, 7*): labile (an aggregate of the three functionally labile classes) and recalcitrant (semi-labile (SL) and semi-refractory (SR) classes).

**Fig. 2.**
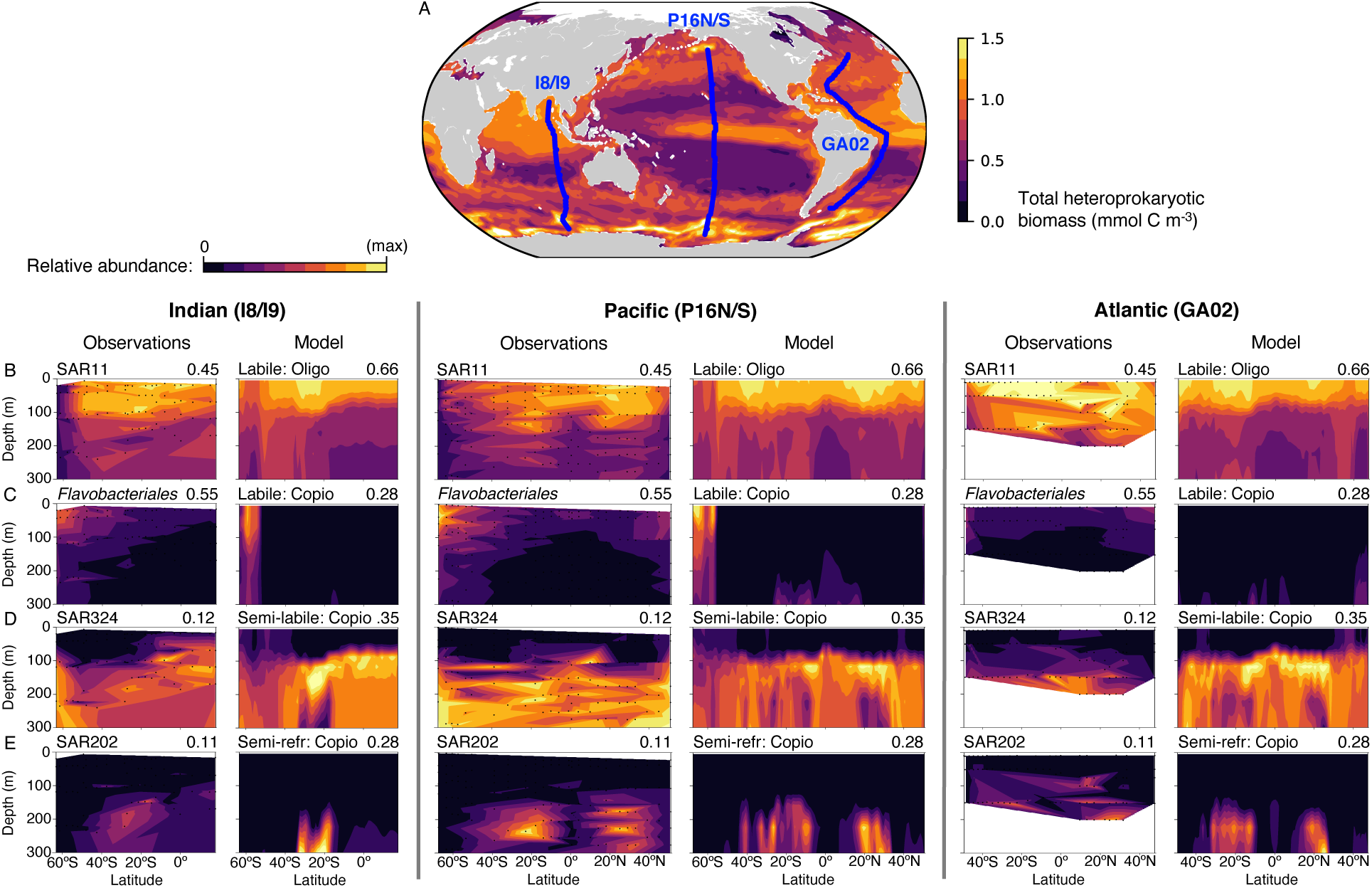
Observed and modeled heteroprokaryotic biogeographies across the three ocean transects. (**A**) The three transects (blue lines) overlaid on the modeled total surface heteroprokaryotic biomass. (**B–E**) Relative abundances of four of the 21 heteroprokaryotic taxonomic guilds and analogous modeled functional types, with the maximum value of the corresponding colorbar noted on each panel. See Fig. S14 for comparable groups of guilds and functional types for which the relative abundance scale is the same. Relative abundance is with respect to non-photosynthetic prokaryotes for ASVs (observations) and functional types (model). We illustrate biogeographies to 300 m to emphasize the strongest vertical gradients; see Figs. S2–S7, S13, and S14 for resolution to 1000 m.

### Global-scale heteroprokaryotic biogeography

We generated a comprehensive, coarse-grained biogeography of the marine heteroprokaryotic metacommunity by leveraging an *in situ* metabarcoding dataset from seawater along latitudinal transects through the Pacific, Indian, and Atlantic ocean basins (Fig. 2A). Our samples spanned the whole heteroprokaryotic community because the samples were not fractionated into size categories. The Shannon diversity index of the amplicon sequence variants (ASVs; unique genetic barcodes) increased from surface waters (roughly 0-150 m) to deeper waters in all transects, whether considering the entire biological community (the complete SSU 16S and 18S rRNA amplicon datasets) or the heteroprokaryotic subset (Fig. S1, Materials and Methods). To reduce the complexity of the datasets, we aggregated the ASVs of all non-phototrophic prokaryotes into ecologically meaningful taxonomic guilds: 21 heteroprokaryotic guilds and two chemoautotrophic (ammonia- and nitrite-oxidizing) guilds (Materials and Methods; Figs. S2–S7; Table S1). The 23 guilds cumulatively represented more than 90% of the non-photoautotrophic prokaryotic abundance (Fig. S8). We aggregated predominantly at the order level or higher, where it has been demonstrated that taxonomy broadly corresponds to function (*26, 27*). We acknowledge that much functional diversity remains unresolved within each guild. We aimed to identify and understand the patterns that manifest at this high level, akin to understanding the large-scale biogeography of trees versus herbaceous plants, despite the enormous functional diversity within these categories.

The biogeographies of the 21 heteroprokaryotic guilds revealed several reoccurring patterns, which we identified by locations of peak relative abundances (Figs. S2–S6). We examined relative abundances with respect to the non-photosynthetic prokaryotic subset, although these patterns remained robust when relative abundances were calculated with respect to the full community (16S and 18S; Materials and Methods). We found that six ‘surface, low latitude/ubiquitous guilds’ had peak relative abundances in the nutrient-depleted ocean deserts. SAR11 was the most abundant representative of this group (Fig. 2B), and is known for its high abundance and oligotrophic traits: slow growth, small cell size, and a high affinity for DOC (*28, 29*). Three ‘surface, high latitude’ guilds were identified by peak relative abundances in high-latitude, nutrient-rich waters (e.g., *Flavobacteriales*, Fig. 2C). Seven ‘deep, ubiquitous’ guilds had negligible relative abundance in surface waters but were present throughout the subsurface (e.g., SAR324 and SAR202, Fig. 2D and E). One ‘deep, sporadic’ guild was identified (*Enterobacterales*; Fig. S5), which we characterized as the most copiotrophic guild. Four ‘particle-like’ guilds (e.g., Marinimicrobia) were broadly ubiquitous, generally peaking at depth but also present in surface waters (Fig. S6), named as such here because this biogeography was consistent with the emergent organic particle-associated populations in the model. The identification of distinct particle-associated guilds confirmed previous work showing that particle association is phylogenetically conserved and so should emerge at a high taxonomic level (*30*).

To complement manual categorization, we conducted a non-metric multidimensional scaling analysis based on the beta-diversity of the heteroprokaryotic communities (measured as Bray-Curtis dissimilarity distances), paired with an agglomerative clustering approach (Materials and Methods). This analysis identified an optimum of three clusters, for which the emergent axes corresponded to high-latitude surface, low-latitude surface, and deep environments (Fig. 3; Fig. S16C). This indicated that the ‘deep, ubiquitous’, ‘particle-like’, and ‘deep, sporadic’ patterns may be better considered as one large deep-peaking category. The categorization of some guilds was ambiguous (e.g., SUP05 relative abundance is both high-latitude- and depth-intensified), and subsets of guilds (e.g., SAR11 ecotypes (*28*)) may be differently categorized. Nevertheless, the repetitive patterns hinted at the potential for a coarse-grained mechanistic explanation of community structure.

**Fig. 3.**
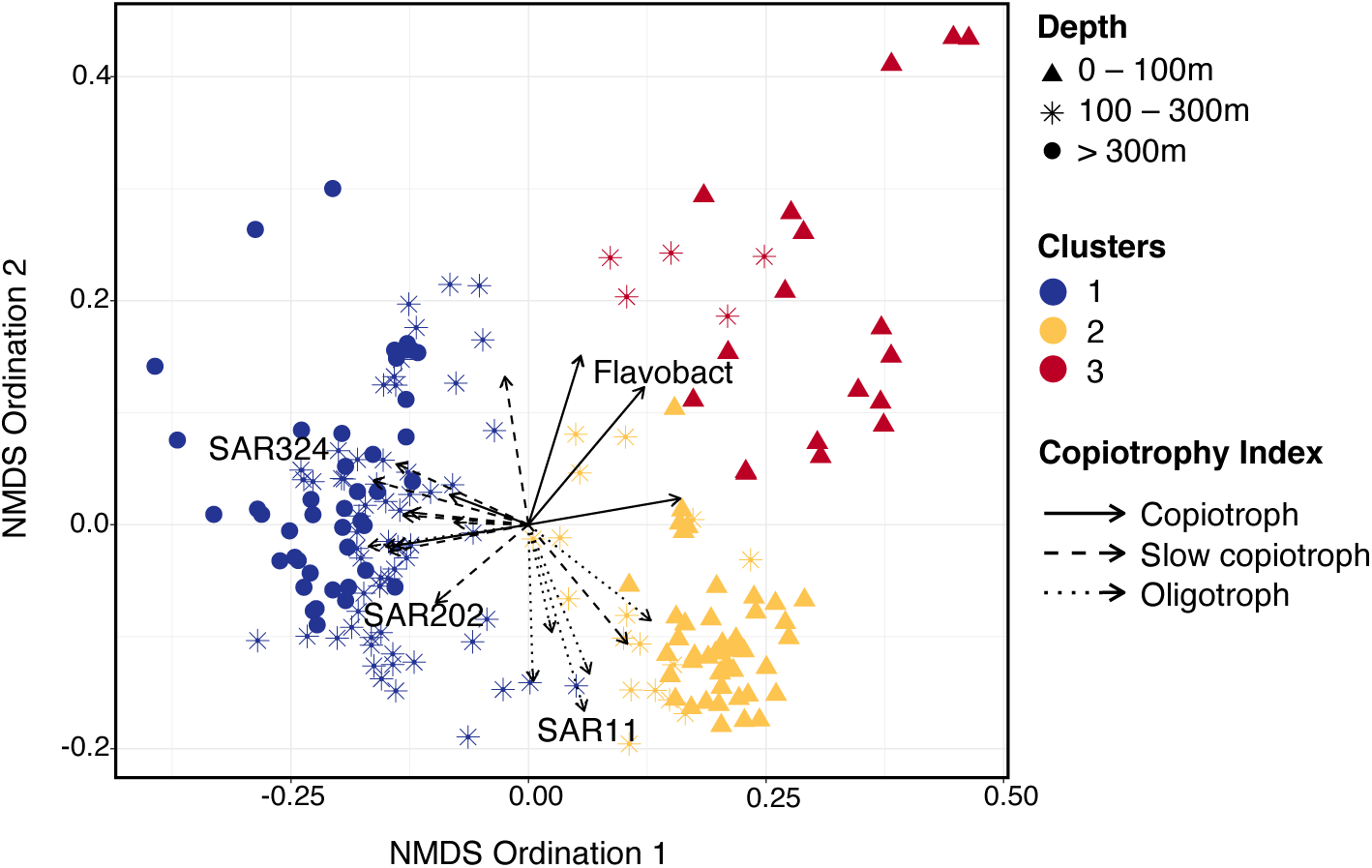
Non-metric multidimensional scaling analysis (NMDS) of the Pacific transect. The axes are determined by analysis of the pairwise similarity among the communities at each sampling location. The symbols indicate the depth of each sample, while the color indicates the results of the hierarchical clustering (Materials and Methods). Arrows for each guild are overlaid in the same ordination space. The direction of each arrow identifies the samples with a general increase in abundance of that guild, and the length of each arrow measures the degree of guild correlation with the ordination axes. See Fig. S16 for annotation with all guild names.

### Modeling heteroprokaryotic functional diversity

We next considered heteroprokaryotic biogeographical patterns from simulations of a mechanistic marine ecosystem model with explicit heteroprokaryotes. The global model synthesized the physics, biogeochemistry, and ecosystem of the ocean (Darwin-MITgcm; Materials and Methods). The ecosystem component allowed for the self-assembly of microbial communities resulting from the interactions among dynamic microbial functional type populations and the environment (*31, 32*). We extended a previous configuration in which all remineralization is due to the growth and respiration of heteroprokaryotic functional types that consume particulate and dissolved organic matter (POM and DOM), as well as multiple phytoplankton, zooplankton, and other chemoautotrophic and anaerobic metabolic functional types (Materials and Methods; Fig. S9). We here expanded the model to include five lability classes of DOM and associated free-living heteroprokaryotic consumers (Fig. 1). We did not resolve refractory and ultra-refractory DOM with millennial lifetimes (*6*). The lability of each class was defined by setting a baseline maximum growth rate for its consumers (*7*) (Fig. 1B and For each lability class, we resolved the rate-affinity tradeoff axis with two competing consumers: (i) small oligotrophs with maximum growth rates lower than the baseline, and with higher substrate affinity, and (ii) larger copiotrophs with maximum growth rates higher than the baseline, and with lower affinity (*17, 18, 20, 21*) (Fig. 1B, Table S2).

DOM was produced from the mortalities and excretions of all biomasses following a lognormal distribution (Fig. 1C), which represented the average outcome from the multiplicative stochasticity of complex underlying processes (*7, 33*). Consequently, cross-feeding and transformations among DOM classes were resolved implicitly. Modeled grazing was structured by organism size, with a preferred prey size for each grazer (*34, 35*). We assumed that all oligotrophs, regardless of lability preferences, were consumed by the smallest grazer, and that the slightly larger copiotrophs were consumed by the second smallest grazer, though model results remained consistent when assuming one shared grazer for all heteroprokaryotes (Materials and Methods). We integrated the model forward in time to reach a quasi-steady state and examined the emergent microbial community structure. Model solutions were qualitatively consistent with observed estimates of global net primary production, the export flux of organic carbon, the inferred pattern of DOC production as a function of lability, DOC accumulation patterns, surface heteroprokaryotic biomass, and surface heteroprokaryotic growth rates (Figs. S10–S11).

We found that the patterns of the emergent heteroprokaryotic biogeographies in the model were remarkably consistent with those exhibited by the taxonomic guilds (Fig. 2, Figs. S12–S14). In modeled surface waters, the most abundant heteroprokaryotic types were the consumers of labile DOM (Fig. 2B and C). Modeled labile DOM, like measured labile constituents, was depleted to low (pM–nM) concentrations, set by the subsistence concentrations of the consumers (*7, 12*) (Fig. S15). This indicated that the three most labile DOM classes could be characterized as functionally labile (*7*), and so we aggregated them and their consumers, which exhibited nearly identical biogeographical patterns (Fig. S15), to consider one labile DOM class. The model captured the latitudinal transition from oligotrophs to copiotrophs in the surface, which was consistent with previous theory and modeling of both phytoplankton and heteroprokaryotes (*24, 35–37*) (Fig. 2B and C). In contrast, all consumers of DOM that was functionally recalcitrant in the surface (*7*) (hereafter referred to as ‘recalcitrant DOM,’ aligning with previously designated semi-labile and semi-refractory classes (*6*); Fig. S10), were largely excluded from the surface, broadly matching the biogeography of the deep, ubiquitous guilds (Fig. 2D and E, Fig. S13). The modeled labile-DOM-consuming copiotroph also became relatively abundant in deeper waters, matching the biogeography of the deep, sporadic *Enterobacterales* guild (Fig. S14).

Quantitative comparison of mathematical ecosystem models and sequencing data remains challenging (*38*). In addition to the difficulty linking coarse-grained representation of metabolism in models to highly detailed, sparse genomic data, we lack an accurate, common currency between model (i.e., biomass concentrations) and molecular data (i.e., sequence counts). To circumvent these barriers in currency and framework, we subsampled colocalized geographical coordinates and depths across the model analog of the Pacific transect and conducted two analyses. First, we conducted a parallel non-metric multidimensional scaling analysis of the beta-diversity of the modeled communities to compare with that of the empirical (Fig. S16; Materials and Methods). Results showed similar ordination of empirical and modeled communities, with a moderately strong positive correlation in a symmetric Procrustes rotation (0.77, p-value: 0.001; Fig. S17). The results of a Mantel test between the model and molecular Bray-Curtis dissimilarity matrices showed a similarly positive correlation (0.63, p-value: 0.001). Both empirical and modeled communities were characterized by three clusters that corresponded predominantly to high-latitude surface, low-latitude surface, and depth.

Second, we conducted a rank correlation analysis by combining the guilds and the modeled functional types into four comparable groups and quantifying the degree to which the relative abundances of these groups were ordered similarly (Fig. S14, Materials and Methods). The resulting Spearman correlation (i.e., consistent ranking of the community structure) at each sampling location was overall positive, and the model matched the observed dominant group at the majority of the locations (Fig. S18). Spearman correlation was highest just below surface waters, at roughly 100-300 m depth (larger colored dots in Fig. S18). However, even with lower correlation, the model captured the dominant group throughout most of the surface ocean (smaller black dots in Fig. S18), where the communities were dominated by SAR11, which indicated that the model did not always skillfully order the less abundant types in the surface ocean, potentially due to uncertainty in filling low biomass, non-dominant group niches.

### Mechanisms shaping heteroprokaryotic community structure

We diagnosed the mechanisms setting the biogeographic patterns in the model. We found that grazing pressure excluded the consumers of recalcitrant DOM from the surface. In surface waters, high biomass of other similar-sized microbial populations sustained high zooplankton grazing, and so the slow-growing recalcitrant-DOM-consuming populations could not be sustained. This “apparent competition” (*39*) was a robust feature of the model. Results held over a wide range in parameter space, within some plausible constraints (Supplementary Text, Fig. S19, Fig. S20). Experimental observations indicate that SAR11 has evolved strategies to escape grazing (*40*), indicating that grazing is indeed a significant obstacle in surface waters.

Recalcitrant DOM accumulated in modeled surface waters because its slow-growing consumers were excluded (Fig. 5A, Fig. S10, Fig. S12). At depth, below the mixed layer, grazing rates waned along with total biomass (Fig. S11), and the slow-growing populations finally became viable. Notably, the slow-growing copiotrophs, rather than the oligotrophs, dominated the deep consumption of recalcitrant DOM (Fig. S13). This was because concentrations of recalcitrant DOM remained relatively high below the mixed layer. The slightly higher maximum growth rates of the slow-growing copiotrophs allowed them to outcompete their oligotrophic counterparts (Fig. S15). The recalcitrant DOM was eventually depleted further at depth, at which point oligotrophic traits were selected. However, at those depths, DOM supply was so low that only low abundances of recalcitrant-DOM-consuming oligotrophs were sustained (Fig. S13) (*41*).

Thus, the model showed how heteroprokaryotic community structure and function can vary with latitude and depth in two ways: (i) a transition in surface waters from the dominance of oligotrophs at low latitudes to the prevalence of copiotrophs at high latitudes, and (ii) a vertical transition from the dominance of oligotrophic populations consuming labile substrates in surface waters to slower-growing, yet copiotrophic populations consuming recalcitrant substrates at depth. The first pattern was driven by the general pattern of increased primary production with latitude, coupled with selection for rate-optimized copiotrophs in dynamic, nutrient-rich waters. The second pattern applied across the lability gradient, and was driven by the alleviation of grazing pressure at depth. The copiotrophs consuming lower quality (recalcitrant) substrates became sustainable at shallower depths relative to their oligotrophic counterparts.

This vertical pattern and its mechanisms were also consistent with previous observations. DOC decreases by about 20–40 μM between the surface and the mesopelagic ocean (*6*), which was captured by the model (Fig. 5A, Fig. S10). Experiments have shown that only microbial communities originating from deep waters can consume the surface-accumulated DOC, even when controlling for the additional nutrients available at depth (*42*). Indeed, time-series data demonstrate that the seasonal downward flux of surface-accumulated DOC is followed by the growth of mesopelagic microbial populations (*43*). It has remained an open question why such mesopelagic populations are not active in surface waters, where their preferred substrates accumulate. The model results here propose high surface grazing rates as one explanation for this phenomenon.

### “Slow copiotrophs” in the deep ocean

The modeled biogeographies of recalcitrant DOM consumers qualitatively matched the biogeographies and putative functions of the SAR324 and SAR202 guilds, respectively (Fig. 2D and E). Thus, the model generated a hypothesis that SAR202 and SAR324, as well as other deep, ubiquitous guilds, may be better classified as copiotrophs than oligotrophs. Specifically, the hypothesis was that these guilds are optimized for relatively fast growth, despite lower absolute maximum growth rates than other guilds due to their subsistence on recalcitrant substrates.

Indeed, sequencing indicates that members of the SAR202 guild metabolize recalcitrant substrates (*44–46*), and that both guilds contain high metabolic versatility and diversity, which is consistent with these guilds having distinct functions (*45, 47, 48*). We also diagnosed that the greater ubiquity of the SAR324 model analog relative to the SAR202 model analog was because of its higher maximum growth rate rather than its higher supply of substrate (Fig. S21, Fig. S22), which allowed us to speculate that SAR324 grow more quickly than SAR202.

To test the hypothesized shift from the dominance of oligotrophy to copiotrophy with depth, we developed a genome-based copiotrophy index. We extracted representative genomes with rRNA sequences that matched ASVs at approximately the genus level (95% sequence identity or greater) from the novel, comprehensive Genome Taxonomy Database (*49*) (Materials and Methods). The 6865 genomes extracted covered about 70% of the communities (Fig. S23). Of these genomes, 43% were metagenome-assembled genomes (MAGs) or single-cell amplified genomes (SAGs), of which 80% were from the ocean. We assembled the copiotrophy index as a function of three copiotrophic characteristics (*19*): (i) the number of genes, (ii) the number of carbohydrate active enzymes (CAZy database) (*50*), and (iii) codon usage bias (Materials and Methods). Codon usage bias is the biased use of alternative codons coding for a particular amino acid that indicates optimization for rapid translation, which, in genes coding for ribosomal proteins, correlates with higher maximum growth rates (*19*). We also used codon usage bias to estimate maximum growth rates for each genome, with and without the confounding impact of temperature (Materials and Methods; Fig. S24). We opted for genus-level matches because our analysis focused on traits known to be consistent across species and strains at this level (*19, 51*), and thus likely more fundamental to determining broad ecological niches (*52*). We demonstrated this consistency with analysis showing minimal variation of the copiotrophy index within species, within genera, and when using 99% sequence identity similarity (Fig. S25, Fig. S26).

For the majority of the guilds, the resulting copiotrophy index values were relatively tightly distributed (Fig. 4), indicating major divisions in growth strategy over evolutionary time, although some guilds (e.g., *Flavobacteriales*) spanned a wide range. A Gaussian mixture analysis showed that the genomes fell into two statistically significant lognormal distributions (Fig. 4, Fig. S25). We labeled these as the canonical “oligotroph” and “copiotroph” clusters, in an update to the classification in previous work (*19*). The oligotroph cluster contained predominantly the surface, low latitude/ubiquitous guilds, including SAR11. The copiotroph cluster contained all of the surface, high latitude guilds as well as *Enterobacterales*. All seven of the deep, ubiquitous guilds fell between these two extremes, and did not clearly belong to either cluster. These deep guilds had some of the lowest maximum growth rate estimates among the guilds (Fig. 4B), even when controlling for temperature (Fig. S27). This indicated that SAR11 and other surface guilds were more oligotrophic but able to grow faster than these deep guilds, decoupling copiotrophy from maximum growth rate (Fig. 4B) (*53*). We labeled these in-between guilds as ‘slow copiotrophs’.

**Fig. 4.**
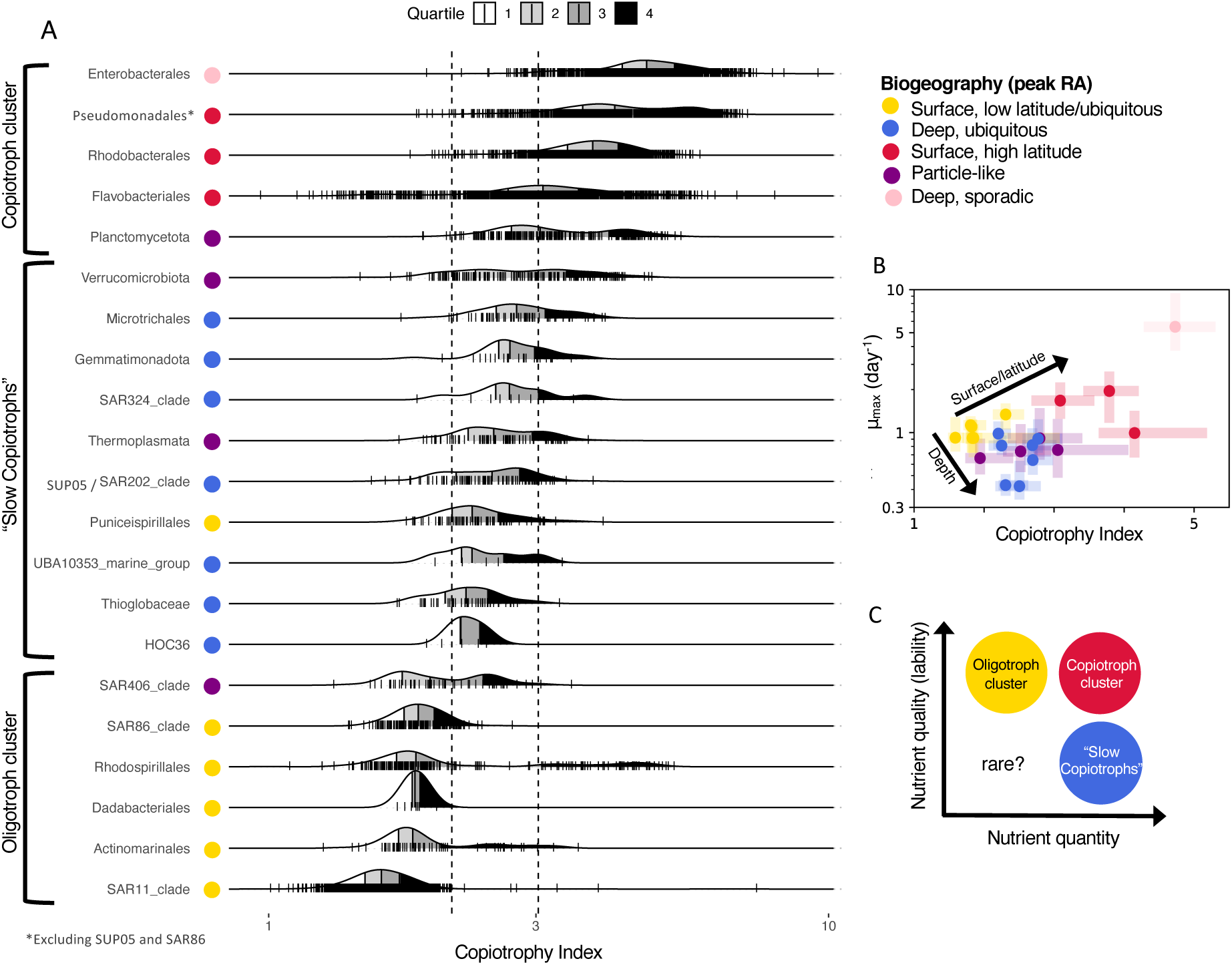
Biogeographical and functional patterns of marine heteroprokaryotic guilds. (**A**) Guilds ranked by the median value of the genome-based copiotrophy index. Dashed lines indicate uncertainty in the clustering analysis, delineating the “slow copiotrophs.” (**B**) Genome-based maximum growth rate (*μ_max_*) estimates (*19*) against the copiotrophy index for each guild (median and interquartile range), colored by biogeography. (**C**) Schematic of proposed broad guild classifications.

At the community level, both model and data showed that the degree of copiotrophy increased with depth as the community-averaged maximum growth rate decreased (Fig. S28, Fig. S29), demonstrating the geography of the decoupling of copiotrophy from maximum growth rate.

Because the sequencing data did not inform the model, the model-data consistency provided two independent lines of evidence for these patterns. In the model, these patterns resulted from the consumption of accumulated (high quantity), recalcitrant (low quality) substrates that did not allow for fast growth in absolute terms (Fig. 4C).

The model reproduced the observed broad pattern of increased microbial diversity in deep waters relative to surface waters (Fig. S29). In the model, slow-growing functional types were excluded from the surface, but few functional types were excluded at depth because continued microbial cycling supplied fresh labile DOM at low rates there, supporting low abundances of many functional types. Therefore, the model proposed higher surface grazing rates as a mechanism for the vertical diversity gradient.

### Ecological controls on ocean carbon cycling and storage

The model allowed us to connect microbial functional diversity to diversity in the rates and locations of organic carbon respiration (Fig. 5). Globally, the labile-DOM-consuming oligotroph – the model analog of SAR11 and other surface, low-latitude/ubiquitous guilds – was responsible for more than 90% of total DOC respiration in surface waters (Fig. 5C). At depth, more diverse communities contributed to DOC respiration, with significant contributions by slow copiotrophs (30–50% of DOC respiration at 200 m), as well as slow oligotrophs (2–20%). While the absolute values of these contributions were not well constrained by the model due to uncertainties in parameters, such as biomass growth efficiencies, the model captured the relative shifts in communities across different biogeochemical regimes and resulting respiration patterns. Therefore, we used the model to evaluate how shifts in ecological dynamics impact carbon cycling and storage.

**Fig. 5.**
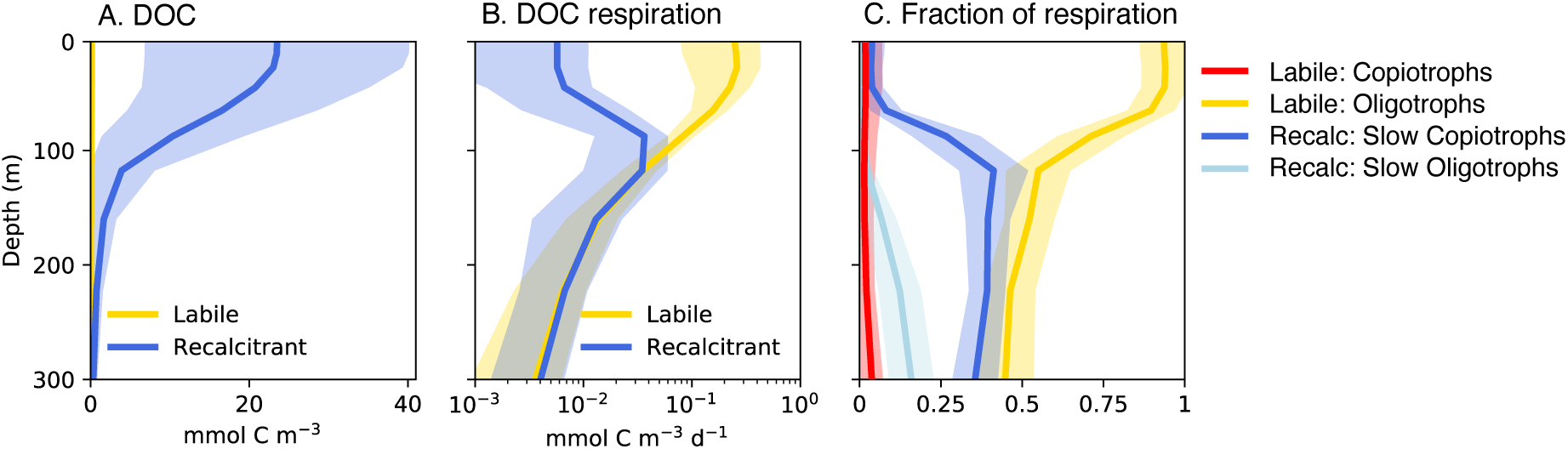
DOC respiration by diverse heteroprokaryotic functional types in the global model. All quantities are modeled global averages, with the shaded area indicating the (volume-weighted) standard deviation. (**A**) Concentrations of labile and recalcitrant DOC. The recalcitrant pool represents the sum of the modeled semi-labile and semi-refractory pools (Fig. 1), neglecting the background refractory concentration of about 40 mmol C m^-3^. (**B**) DOC respiration rates. (**C**) Fraction of respiration by heteroprokaryotic functional types.

Because labile DOC respiration was predominantly limited by substrate availability, and thus globally dominated by high-affinity oligotrophs, changes to labile DOC supply were closely matched by concurrent changes in respiration, with little impact on DOC storage. Furthermore, the geography of labile DOC cycling was robust to changes in grazing dynamics (Fig. S20). In contrast, recalcitrant DOC accumulated in the surface because grazers suppressed its slow consumers, and thus, in a model simulation without explicit grazers, surface DOC was reduced by 90% because the slow recalcitrant DOC consumers persisted there (Fig. S20C). Without apparent competition from grazers, the slow consumers’ growth was sufficient to match mortality rates, enabling recalcitrant DOC consumption in the surface. Therefore, the degree of recalcitrant DOC accumulation – the bulk of modeled DOC storage – was sensitive to the balance of growth and mortality of its consumers. The model experiment demonstrated that relatively small changes in this balance, when sufficient to allow consumers to sustain their populations in surface waters, corresponded to a relatively large decrease in DOC storage.

### Outlook

Our results demonstrated an “emergent simplicity” (*27*) and qualitative, coarse-grained predictability of the complex marine microbiome at the global scale. Resolving heteroprokaryotic diversity with two axes – organic substrate lability and a rate-affinity tradeoff – was sufficient to broadly capture observed biogeographical patterns and simple enough to understand the mechanisms producing them. The fact that the model demonstrated the same broad patterns as the ASV and genomic data, without the input of these data to the model, bolstered our conclusions and provided a mechanistic explanation for the patterns. Results linked microbial diversity to biogeochemical function and proposed new heteroprokaryotic functional types: slow copiotrophs. Incorporating other traits and tradeoffs into the model would resolve additional functional diversity (*54–59*).

By linking microbial growth to the turnover of organic carbon, our model results confirm that ecological interactions matter for ocean DOC storage (*7, 8*). This challenges ocean biogeochemical models having a simpler resolution of respiration to mechanistically capture the controls on DOC accumulation and potential feedbacks to warming. Our model showed that if ecological or biochemical dynamics shifted such that slow copiotrophs could persist in the surface, standing stocks of surface DOC would be reduced.

We have identified the mechanisms shaping the coarse-grained structure and function of marine heteroprokaryotic communities in the ocean. In doing so, we have made progress towards meeting two grand challenges in marine biogeochemistry: organizing the functional diversity of complex microbial communities along quantitative axes, and connecting sequencing datasets with ecosystem models. Additionally, the model represents a strategy for incorporating dynamic heteroprokaryotic feedbacks into earth system models. This strategy can be used to investigate heterogeneity in the response of microbial communities impacting biological carbon storage in a warmer climate.

### Methods

Please see the attached Supporting Information for Methods, supplemental figures, and supplemental tables. Note that all references (including those for the Methods) are listed here in the main text.

## Acknowledgments

We thank Nathan Williams for help with sequencing data access. We thank Steven Biller and Paul Berube for providing DNA to generate amplicons from the GEOTRACES GA02 cruise. CAC thanks the US GO-SHIP program and the US National Science Foundation for support in collection of samples on P16N/S and I8/I9 cruises. Funding: Simons Foundation Postdoctoral Fellowship in Marine Microbial Ecology (EJZ, JLW). Carnegie Institution for Science (EJZ, LX). National Science Foundation grant 2125142 (EJZ, JAF). Simons Foundation grant 549943FY22 (JAF). Simons Collaboration on Computational Biogeochemical Modeling of Marine Ecosystem/CBIOMES grant 549931 (SD). Simons Foundation International BIOS-SCOPE program (CAC). Simons Foundation grant 542389 (NML).

## Data availability

All data and model code are freely available: processed amplicon data (*66*), ASV-to-genome matching and taxonomic analysis (*87*), code for copiotrophy index analysis (*88*), and Darwin-MITgcm model code (*89*).

## References

1. P. G. Falkowski, T. Fenchel, E. F. Delong, Science 320, 1034 (2008).

2. T. Volk, M. I. Hoffert, The carbon cycle and atmospheric CO_2_: Natural variations Archean to present. Chapman conference papers, 1984, E. T. Sundquist, W. S. Broecker, eds. (American Geophysical Union, 1985), pp. 99–110.

3. R. Séférian, et al., Current Climate Change Reports 6, 95 (2020).

4. W. R. Wieder, G. B. Bonan, S. D. Allison, Nature Climate Change 3, 909 (2013).

5. S. A. Henson, et al., Nature Geoscience 15, 248 (2022).

6. D. A. Hansell, Annual Review of Marine Science 5, 421 (2013).

7. E. J. Zakem, B. B. Cael, N. M. Levine, Proceedings of the National Academy of Sciences 118, e2016896118 (2021).

8. S. T. Lennartz, D. P. Keller, A. Oschlies, B. Blasius, T. Dittmar, Global Biogeochemical Cycles 38, e2023GB007912 (2024).

9. J. I. Hedges, et al., Organic Geochemistry 31, 945 (2000).

10. C. A. Carlson, D. A. Hansell, Biogeochemistry of Marine Dissolved Organic Matter (Elsevier Inc., 2015), chap. 3, pp. 65–126.

11. A. D. Steen, et al., ISME Journal 13, 3126 (2019).

12. M. A. Moran, et al., Nature Microbiology 7, 508 (2022).

13. S. Sunagawa, et al., Science 348, 1261359 (2015).

14. V. J. Coles, et al., Science 358, 1149 (2017).

15. F. Milke, J. Meyerjürgens, M. Simon, Nature Communications 14, 6141 (2023).

16. J. T. Lennon, et al., mBio 15, e00455 (2024).

17. A. L. Koch, BioEssays 23, 657 (2001).

18. F. M. Lauro, et al., Proceedings of the National Academy of Sciences 106, 15527 (2009).

19. J. L. Weissman, S. Hou, J. A. Fuhrman, Proceedings of the National Academy of Sciences 118, e2016810118 (2021).

20. A. Soler-Bistué, L. L. Couso, I. E. Sanchez, Environmental Microbiology 25, 1232 (2023).

21. N. Norris, N. M. Levine, V. I. Fernandez, R. Stocker, PLoS Computational Biology 17, e1009023 (2021).

22. M. Wietz, L. Gram, B. Jørgensen, A. Schramm, Aquatic Microbial Ecology 61, 179 (2010).

23. S. Yooseph, et al., Nature 468, 60 (2010).

24. M. V. Brown, M. Ostrowski, J. J. Grzymski, F. M. Lauro, Marine Genomics 15, 17 (2014).

25. C. Carlson, H. Ducklow, Deep Sea Research Part II: Topical Studies in Oceanography 42, 639 (1995).

26. S. Louca, L. W. Parfrey, M. Doebeli, Science 353, 1272 (2016).

27. J. E. Goldford, et al., Science 361, 469 (2018).

28. S. J. Giovannoni, Annual Review of Marine Science 9, 231 (2017).

29. S. E. Noell, S. J. Giovannoni, Environmental Microbiology 21, 2559 (2019).

30. G. Salazar, et al., Molecular Ecology 24, 5692 (2015).

31. M. J. Follows, S. Dutkiewicz, S. Grant, S. W. Chisholm, Science 315, 1843 (2007).

32. S. Dutkiewicz, et al., Nature Climate Change 5, 1002 (2015).

33. D. C. Forney, D. H. Rothman, Journal of The Royal Society Interface 9, 2255 (2012).

34. B. A. Ward, S. Dutkiewicz, O. Jahn, M. J. Follows, Limnology and Oceanography 57, 1877 (2012).

35. C. L. Follett, et al., Proceedings of the National Academy of Sciences 119, e2110993118 (2022).

36. S. Dutkiewicz, M. J. Follows, J. G. Bragg, Global Biogeochemical Cycles 23, GB4017 (2009).

37. C. I. Abreu, M. D. Bello, C. Bunse, J. Pinhassi, J. Gore, Science Advances 9, eade8352 (2023).

38. S. Meiler, et al., Limnology and Oceanography 67, 816 (2022).

39. R. D. Holt, Theoretical Population Biology 12, 197 (1977).

40. A. Dadon-Pilosof, et al., Nature Microbiology 2, 1608 (2017).

41. M. L. Sogin, et al., Proceedings of the National Academy of Sciences 103, 12115 (2006).

42. C. A. Carlson, et al., Limnology and Oceanography 49, 1073 (2004).

43. S. Liu, et al., Frontiers in Microbiology 13, 833252 (2022).

44. Z. Landry, B. K. Swa, G. J. Herndl, R. Stepanauskas, S. J. Giovannoni, mBio 8, 00413 (2017).

45. J. H. Saw, et al., mBio 11, 02975 (2020).

46. S. Liu, et al., Limnology and Oceanography 65, 1532 (2020).

47. C. S. Sheik, S. Jain, G. J. Dick, Environmental Microbiology 16, 304 (2014).

48. D. Boeuf, et al., Microbiome 9, 172 (2021).

49. D. H. Parks, et al., Nucleic Acids Research 50, D785 (2022).

50. E. Drula, et al., Nucleic Acids Research 50, D571 (2022).

51. S. Vieira-Silva, E. P. C. Rocha, PLoS Genetics 6, e1000808 (2010).

52. A. C. Martiny, K. Treseder, G. Pusch, The ISME Journal 7, 830 (2013).

53. L. L. Couso, A. Soler-Bistué, A. A. Aptekmann, I. E. Sánchez, Environmental Microbiology 25, 3052 (2023).

54. T. F. Thingstad, R. Lignell, Aquatic Microbial Ecology 13, 19 (1997).

55. J. L. Green, B. J. M. Bohannan, R. J. Whitaker, Science 320, 1039 (2008).

56. S. D. Allison, Ecology Letters 15, 1058 (2012).

57. J. B. Martiny, S. E. Jones, J. T. Lennon, A. C. Martiny, Science 350 (2015).

58. E. Litchman, K. F. Edwards, C. A. Klausmeier, Frontiers in Microbiology 6, 254 (2015).

59. M. Gralka, S. Pollak, O. X. Cordero, Nature Microbiology 8, 1799 (2023).

60. C. M. Swan, D. A. Siegel, N. B. Nelson, C. A. Carlson, E. Nasir, Deep Sea Research Part I: Oceanographic Research Papers 56, 2175 (2009).

61. R. F. Anderson, Annual Review of Marine Science 12, 49 (2020).

62. S. J. Biller, et al., Scientific Data 5, 180176 (2018).

63. K. H. Boström, K.Simu, Å. Hagström, L.Riemann, Limnology and Oceanography: Methods 2, 365 (2004).

64. M. Manganelli, et al., PloS one 4, e6941 (2009).

65. C. N. Signori, F. Thomas, A. Enrich-Prast, R. C. Pollery, S. M. Sievert, Frontiers in microbiology 5, 647 (2014).

66. McNichol, J., Processed ASV tables and associated information from the Global rRNA Universal Metabarcoding of Plankton (GRUMP) Project, osf.io/57dpa/ (2024).

67. A. E. Parada, D. M. Needham, J. A. Fuhrman, Environmental Microbiology 18, 1403 (2016).

68. C. Quast, et al., Nucleic Acids Research 6, D590 (2013).

69. L. Guillou, et al., Nucleic Acids Research 41, D597 (2012).

70. J. McNichol, P. M. Berube, S. J. Biller, J. A. Fuhrman, Msystems 6, 10 (2021).

71. J. D. Hunter, Computing in Science & Engineering 9, 90 (2007).

72. P. Yarza, et al., Nature Reviews Microbiology 12, 635 (2014).

73. J. Zheng, Q. Ge, Y. Yan, X. Zhang, L. Huang, Nucleic Acids Research 51, W115 (2023).

74. R Core Team, R: A Language and Environment for Statistical Computing, R Foundation for Statistical Computing, Vienna, Austria (2022).

75. L. Scrucca, M. Fop, T. B. Murphy, A. E. Raftery, The R Journal 8, 289 (2016). 76.

76. J. Weissman, et al., bioRxiv p. 2021.10.15.464604 (2021).

77. T. Seemann, Bioinformatics 30, 2068 (2014).

78. C. Wunsch, P. Heimbach, Physica D 230, 197 (2007).

79. E. J. Zakem, et al., Nature Communications 9, 1206 (2018).

80. E. J. Zakem, N. M. Levine, Global Biogeochemical Cycles 33, 1389 (2019).

81. L. A. Anderson, Deep Sea Research Part I 42, 1675 (1995).

82. E. J. Zakem, M. J. Follows, Limnology and Oceanography 62, 795 (2016).

83. E. J. Zakem, M. F. Polz, M. J. Follows, Nature Communications 11, 5680 (2020).

84. E. Litchman, C. A. Klausmeier, O. M. Schofield, P. G. Falkowski, Ecology Letters 10, 1170 (2007).

85. J. Guo, et al., Geophysical Research Letters 50, e2023GL102896 (2023).

86. S. Dutkiewicz, et al., Biogeosciences 12, 4447 (2015).

87. McNichol, J., github.com/jcmcnch/MicroheterotrophModelling, doi:10.6084/m9.figshare.28236062 (2025).

88. Weissman, J. L., github.com/jlw-ecoevo/bhet_code, doi:10.6084/m9.figshare.28235984 (2025).

89. Zakem, E. J., github.com/emilyzakem/heteroprokaryotes, doi:10.5281/zenodo.14681401 (2025).

90. J. H. Martin, G. A. Knauer, D. M. Karl, W. W. Broenkow, Deep Sea Research Part A. Oceanographic Research Papers 34, 267 (1987).

91. D. L. Kirchman, Annual Review of Marine Science 8, 285 (2016).

92. R. Schlitzer, Geophysical Monograph 114, 107 (2000).

93. S. A. Henson, et al., Geophysical Research Letters 38, L04606 (2011).

94. D. Siegel, et al., Global Biogeochemical Cycles 28, 181 (2014).

95. H. Ducklow, Microbial Ecology of the Oceans, D. L. Kirchman, ed. (Wiley-Liss, Inc, 2000), chap. 4, pp. 85–120.

96. A. W. Omta, E. A. Heiny, H. Rajakaruna, D. Talmy, M. J. Follows, Ecological Modelling 475, 110183 (2023).

