## Supporting Information for "Functional biogeography of marine microbial heterotrophs"

*Note: See main text for the numbered list of references.*

### Methods

#### S1 Data analysis

#### S1.1 Sequencing data analysis

Microbial distributions were determined from 3-domain metabarcoding of unfractionated ( > 0.2 μm) seawater samples. Samples from repeat hydrography lines P16S/P16N and I8N/I8S were collected in 2005/2006 and 2007, respectively, as part of the GO-SHIP (CLIVAR) program (*60*). Samples from cruise GA02 were collected in 2010/2011 as part of the GEOTRACES program (*61*). GA02 DNA was the same material used for BioGEOTRACES metagenomes (*62*). For GO-SHIP samples, filters were stored at -80°C in lysis buffer until DNA extraction was performed in 2020-2021. DNA was extracted as previously described (*63, 64, 65*) with the addition of a bead-beating step (*66*). Metabarcodes were generated using standard protocols (see: https://www.protocols.io/view/fuhrman-lab-515f-926r-16s-and-18s-rrna-gene-sequen-j8nlkpd1g5r7/v2) and the 515Y/926R primer set (*67*). Bioinformatic analysis was conducted with qiime2-2022.2, using DADA2 for denoising following a workflow specific for this primer set: https://github.com/jcmcnch/eASV-pipeline-for-515Y-926R. Amplicon sequence variants were assigned taxonomy with SILVA138.1 and PR2 4.14.0 (*68, 69*). Processed amplicon data is available at the Open Science Foundation (*66*): https://osf.io/57dpa/. In order to generate a table containing only the organisms of interest, 18S sequences, Cyanobacteria, and organellar 16S (i.e., plastid/mitochodria) were excluded using unique taxonomic identifiers. This final ASV table retained only non-photosynthetic prokaryotes: heterotrophic Bacteria/Archaea and chemoautotrophic taxa (e.g., *Nitrospinales* and *Nitrosopumilales*). Since water samples were not size-fractionated, this community characterization included both "free-living" and "particle-associated" taxa, providing a holistic, comprehensive picture of the microbial community (*70*).

**S1.2 Taxonomic guilds**

A manually curated list of 21 heteroprokaryotic "guilds" was developed based on SILVA 138.1 taxonomy (*68*). These guilds represent dominant heterotrophic organisms on the three ocean transects. Thirteen of the guild names mapped to the order level (e.g., *Rhodobacterales*, SAR11, *Flavobacteriales*), six mapped to coarser taxonomies or poorly-annotated groups (e.g., *Verrucomicrobiota*, UBA10353 marine group), and two were split from the order Pseudomonadales due to their known unique biogeographies (SAR86 and *Thioglobaceae* / SUP05). All bioinformatic steps were recorded in a github repository: https://github.com/jcmcnch/MicroheterotrophModelling.

Fig. S8 shows coverage of the community by the 21 heteroprokaryotic guilds and the two nitrifying guilds (using keywords Nitroso* and Nitrosp* for ammonia-oxidizing and nitrite-oxidizing populations, respectively), both with respect to the full community (all 16S and 18S ASVs) and the non-photosynthetic prokaryotic subset. Figs. S2–S7 show the biogeographies of each of the 23 guilds, grouped into the broad biogeographic patterns. Each plot shows the summed relative abundance of the ASVs within each guild relative to the non-photosynthetic prokaryotic subset of the data. (See S1.3 for details.) We interpolated the data using the 'griddata' function from Python's SciPy library, using the linear method of interpolation. The colorbars vary among the different guilds (though are the same for each guild in all three transects) to emphasize the biogeographical pattern for each guild (we allowed Matplotlib (*71*) to choose the colorbar range to be as objective as possible). Plots with relative abundance (RA) with respect to the full community (all 16S and 18S ASVs), are available in a folder named "GuildRA_16Sand18S" here: https://github.com/emilyzakem/heteroprokaryotes.

### S1.3 Calculations of relative abundances

We used relative abundance (RA) for qualitative comparisons of heteroprokaryotic biogeography and patterns between model and observations. This was because it was not possible to more directly connect the modeled biomass concentrations with sequence counts, and also because we did not necessarily expect biomass of the modeled functional groups to directly align with that of the taxonomic guilds, since we considered the functional groups to largely overlap with, but not exactly correspond to, taxonomy. For the observed guilds (Fig. 2, Figs. S2–S7), the RA was calculated by summing up the relative abundances of each of the ASVs within each guild, with the denominator as the total abundance of the non-photosynthetic prokaryotic abundance at each sampling location. For the model (Fig. 2, Fig. S13), the RA was calculated conceptually in the same way, as the ratio of resulting biomass of the functional type to the total biomass of all non-photosynthesizing, non-zooplankton functional types. We needed to compare the RA of the guilds to the modeled types in this way, with the non-photosynthetic prokaryotic total in the denominator, to enable an 'apples-to-apples' comparison between model and data. We also generated plots of the RA of the guilds with respect to the full community (i.e., using the full 16S and 18S in the denominator) in a folder named "GuildRA_16Sand18S" located here: https://github.com/emilyzakem/heteroprokaryotes.

The biogeographical patterns remained the same although the magnitude of the relative abundances decreased (primarily in the surface, where photosynthetic prokaryotic and eukaryotic abundance is higher; Fig. S8) due to the larger denominator. For the Shannon diversity calculations in Fig. S1A, the relative abundances were calculated with respect to the entire 16S and 18S dataset.

**S1.4 Copiotrophy Index**

To enable specific functional insights into the guilds, BLASTn was used to map ASV sequences to representative SSU rRNA sequences from version r220 of the Genome Taxonomy Database (GTDB) (*49*) at approximately genus-level resolution (95% sequence identity) (*72*). This allowed us to directly associate ASV sequences with genomic information from close relatives. GTDB draws on all metagenome-assembled genomes (MAGs) or single-cell amplified genomes (SAGs) that have been uploaded to NCBI, with additional filtering for quality. Of the 6865 genomes used in the current analysis, 2950 were MAGs or SAGs, and at least 2353 were MAGs or SAGs from the ocean (annotated as coming marine, ocean, deep-sea, cold seep, or saline water samples in the metadata, with many genomes lacking a source annotation, making this a lower bound). Genomic insights into these organisms could then mapped back onto the ASV relative abundances across the transects to visualize broad patterns and intercompare with modeling results. All bioinformatic steps were recorded in a github repository: https://github.com/jcmcnch/MicroheterotrophModelling.

We calculated the copiotrophy index on the P16 transect (the most globally extensive transect capturing the largest range in ecotypes) as follows. We annotated the carbohydrate active enzymes in each genome in our ASV-matched dataset using dbCAN3 on each set of annotated proteins available from GTDB220 (default protein settings, retaining proteins annotated by at least two of dbCAN's internal methods) (*73*). For each genome in our ASV-matched dataset, we performed principal component analysis (prcomp function with scaling from the stats R package) (*74*) on three variables: (i) the number of carbohydrate active enzymes output by dbCAN, (ii) the total number of open reading frames annotated by prokka, and (iii) normalized codon useage bias (dCUB) as output by gRodon (see next section, "S1.5 Maximum growth rate estimates," for more details). Each of these three traits is often individually associated with copiotrophy (*19*). We then took the first principle component after this transformation and transformed this combined value to be greater than one by negating it (because more negative dCUB values correspond to a faster growth rate), and adding its minimum value to one. Thus larger values of this index corresponded to a greater degree of copiotrophy according to genomic indicators. Clustering to assess separation along this axis was performed using a gaussian mixture model (Mclust function from mclust R package (*75*)), with a "confident" classification considered where there was 90% certainty in the classification, and all other classifications considered to be uncertain (Fig. S25). We then applied the same annotation approach to the two other transects (I8/I9 and GA02) by applying the PCA weightings and classification thresholds calculated from P16. All code for this analysis has been documented: https://github.com/jlw-ecoevo/bhet_code.

We calculated the weighted average of the copiotrophy index along the transect in three different ways, all resulting in a similar qualitative pattern (Fig. S28, with details in the caption). In both Fig. S28C and Fig. S29A, the copiotrophy index values for each guild were discretized in order to compare most directly with the global model by assigning a value of 0 to oligotrophs and 1 to slow copiotrophs and copiotrophs (the `fraction of copiotrophy'). This matched the model configuration, where functional types were considered either copiotrophic or oligotrophic, allowing for the closest 'apples-to-apples' comparison between model and data. Thus, the modeled fraction of copiotrophy in Fig. S29D was calculated analogously to the data by assigning the oligotrophic types a value of 0 and the copiotrophic types (including the particle-associated consumers) a value of 1, and then calculating the average weighted by the biomasses of each type.

#### S1.5 Maximum growth rate estimates

We used gRodon2 to estimate maximum growth rates from codon usage statistics (*19*). Out of the 60,524 ASVs across the three transects (all non-photosynthetic prokaryotes, which includes nitrifying microorganisms), 33,405 (55%) were matched with at least one genome that had at least 10 ribosomal proteins, allowing for an estimate of the maximum growth rate for that ASV. These ASVs tended to be more abundant, with the median relative abundance per sample of genome-matched ASVs ranging from 69%–70% per transect (Fig. S23). For each ASV, we calculated an optimal growth temperature with respect to each transect by taking the top 1% of samples in terms of that ASV's relative abundance in that transect and calculated the mean temperature recorded for those samples. This approach has been applied previously to estimate the optimal growth temperature of MAGs with similar performance to more complex machine-learning models (*76*). We then took all genomic hits from GTDB (*49*) for each ASV and fed each into gRodon2 with the corresponding optimal growth temperature estimate (in "full" prediction mode with the "madin" training set (*19*)) after annotating coding sequence and ribosomal proteins with prokka (*77*) (with options –norrna –notrna –centre X –compliant, and –kingdom

Archaea when the organism was an archaeon). Finally, we took the average across genomic hits in GTDB for each ASV to obtain both the temperature-corrected minimum doubling time for each ASV as well as an estimate of the normalized codon usage bias (dCUB). We also ran the same estimation procedure assuming all ASVs had an optimal growth temperature of 20°C to compare across ASVs growing at different temperature ranges.

In ref. (*19*), a five-hour doubling time was discussed as the cutoff for reliable predictions. This cutoff was not appropriate for our estimates for two reasons. First, this cutoff corresponded to a particular choice of optimum temperature for the estimates, and so was not appropriate for our estimates that assume significantly lower optimum temperatures (either 20°C or the temperature associated with the highest relative abundances of each ASV as explained above). Second, more critically, this cutoff derived from a clustering analysis that we have here updated using the copiotrophy index. In the previous work, the 'oligotrophic' cluster was below the cutoff, while the 'copiotrophic' cluster was above the cutoff. Here, our new clustering analysis identified two new clusters, with the addition of many ASVs not clearly falling into one or the other cluster. However, similar to the previous work, we interpreted growth rate estimates for the oligotrophic cluster as highly uncertain, and should be best considered as upper-bound estimates.

Fig. S24A and B shows the frequency of the resulting maximum growth rate estimates for all of the non-photosynthetic prokaryotes with gRodon estimates from the P16 transect (n = 18, 831). Because the slower growth rates are best considered as upper-bound estimates, we expect that the distribution is wider than illustrated, spreading to higher frequencies at lower maximum growth rates. We have illustrated the resulting distributions considering two different estimates of the temperature optimum for each of the ASVs represented: 1) with the temperature optimum as the mean temperature from the samples that comprised that ASV's top 1% relative abundance, as explained above ("Variable $T_{opt}$"), and 2) with the temperature optimum the same ($T_{opt}$ = 20°C) for all. The variable $T_{opt}$ estimate took into account that the organisms may have adapted to low temperatures, but involved much uncertainty since peak relative abundance is controlled by many factors other than temperature. The $T_{opt}$ = 20°C estimate allowed for comparisons of maximum growth rate without any impact of temperature, and critically, a direct comparison with the model maximum growth rate parameter which had a reference temperature of 20°C. However, in reality, many deep ocean types may have adapted to low temperatures and so may not grow optimally at higher temperatures in a laboratory setting. Fig. S24E and F shows the resulting community-averaged (relative-abundance-weighted) maximum growth rates across the Pacific transect, compared to the modeled community averages. When controlling for temperature ($T_{opt}$ = 20°C), the empirical community-averaged maximum growth rate decreased with depth in the subtropical gyres, but not at the equator, where there was fine-scale variability showing both lower and higher rates compared to the surface. It also increased with latitude in the surface, reflecting the shift from dominant oligotrophs at low latitudes to higher abundances of copiotrophs at high latitudes. This was consistent with the modeled patterns.

The weighted average genome-based maximum growth rate ($\mu_{max}$) from the data (Fig. S29B) was calculated by weighting the $\mu_{max}$ estimate by the relative abundance of each ASV by the relative abundance with respect to all ASVs that had genome representatives (and thus $\mu_{max}$ estimates). (Thus, the relative abundance of ASVs without $\mu_{max}$ estimates were not included in the denominator.) For the model analog (Fig. S29E), the average $\mu_{max}$ was calculated as similarly as possible by weighting the $\mu_{max}$ of each non-photosynthetic, non-zooplankton functional type by its biomass concentration with respect to total non-photosynthetic, non-zooplankton biomass.

**S2 Model details and configuration**

The global ocean model (the Darwin-MITgcm model) resolved global biogeochemistry dynamically from the ecological interactions of explicit microbial populations. Specifically, it resolved the cycling of carbon, nitrogen, phosphorus, silica, iron, and oxygen by tracking the concentrations of microbial functional type biomass, inorganic nutrients, and non-living organic nutrients, and was coupled to the ECCO-GODAE state estimate of the ocean circulation ($1° \times 1°$ horizontal resolution; 23 vertical levels) (*31, 32, 78*). The configuration here combined previous introduction of diverse microbial metabolic functional types into the model (*79*) with the rate-affinity tradeoff that was introduced for heteroprokaryotes competing for one bulk class of organic matter (*35*), which built upon the allometric scaling that parameterizes phytoplankton and zooplankton traits. Six phytoplankton types, five zooplankton types, a chemoautotrophic aerobic ammonia-oxidizing functional type and a chemoautotrophic aerobic nitrite-oxidizing functional type were also resolved (*79*). Section S2.1 explains the new addition of the multiple heteroprokaryotic functional types and the organic matter pools that they consume, including the resolution of both aerobic and anaerobic types. Code for this particular Darwin-MITgcm configuration has been made available here: https://github.com/emilyzakem/heteroprokaryotes. Model details, equations, and parameter values for other (non-heteroprokaryotic) functional types, light, and nutrients can be found in the publication of a previous model version (*79*) and at: https://darwin3.readthedocs.io/en/latest/phys_pkgs/darwin.html.

We initialized the model from climatological nutrient concentrations and equal biomass concentrations of all functional types at all locations, and we numerically integrated the model forward in time until a quasi-steady state was reached. All plots illustrate annually averaged model solutions after 100 years of integration. This was a sufficient length of integration to investigate patterns in the top few hundred meters of the model (and definitely to 300 m, as illustrated in the main text). However, we did not run the model long enough to capture the impacts of the overturning circulation at deeper depths, or to resolve the refractory and ultra-refractory DOC pools with millennial lifetimes (*6*).

#### S2.1 Heteroprokaryotic functional types

The model resolved the variation in time of the biomass concentration of each heteroprokaryotic functional type $i$ ($B_i$ in mmol C m$^{-3}$) as:

$$\frac{\partial B_i}{\partial t} = (\mu_i - L_i)B_i - \nabla \cdot (\mathbf{u}B_i) + \nabla \cdot (\boldsymbol{\kappa}\nabla B_i) \quad \text{(S1)}$$

for growth rate $\mu_i$ (d$^{-1}$), biomass loss rate $L_i$ (d$^{-1}$), water velocities **u**, and mixing coefficients **κ**. For the aerobic heteroprokaryotic functional types illustrated and analyzed in the main text, growth rate was determined from the limiting growth rate on organic carbon (C), organic and inorganic nitrogen (N), phosphorus (P), and iron (Fe), or oxygen ($O_2$; though $O_2$ was never limiting to the aerobic types along the three transects), assuming Liebig's law of the minimum, as:

$$\mu_i = \min[\mu_{C_i}, \mu_N, \mu_P, \mu_{Fe}, \mu_{O_2}]\,\gamma_T \quad \text{(S2)}$$

and impacted by the local water temperature by the temperature modification function $\gamma_T$ as described below (Eqn. S5). The stoichiometric relationships between the organic and inorganic elemental pools followed a previous model version (*80*), such that heteroprokaryotes were able to also assimilate inorganic ammonium and phosphate, although (as demonstrated (*80*)), growth was predominantly limited by organic C in the oxygenated domain of the model ocean, since the majority of utilized organic C is oxidized for energy according to the biomass yield $y_C$ (mol B (mol C)$^{-1}$). The organic-C-limited growth rate for functional type $i$ consuming dissolved or particulate organic C pool $j$ was calculated following a Michaelis-Menton saturating function:

$$\mu_{C_i} = y_C V_{max_{OM_i}} \frac{C_j}{C_j + k_{C_i}} \gamma_T \qquad \text{(S3)}$$

where $V_{max_{OM_i}}$ (d$^{-1}$) is the specific maximum uptake rate of organic carbon for each functional type. Thus, the maximum growth rate of each functional type listed in Table S2 was controlled by two parameters: $\mu_{max_{C_i}} = y_C V_{max_{OM_i}}$. However, for simplicity, $y_C$ was set as constant (0.3 mol biomass synthesized per mol organic C consumed) for all functional types. The yield in reality varies among heterogeneous organic substrates and populations, but here, we varied just one of the two parameters impacting $\mu_{max_{C_i}}$ because the model solutions were qualitatively independent of this choice. Limitation by organic and inorganic nitrogen, phosphorus, and iron followed an analogous Michaelis-Menton relationship (*80*). The relative half-saturation constants were set by assumed constant stoichiometry of heteroprokaryotic biomass (*80*): C:N = 5 (lower than phytoplankton), with other biomass elemental ratios (N:P:Fe) equal to those of phytoplankton (*81*; https://darwin3.readthedocs.io/en/latest/phys_pkgs/darwin.html) so that the C:N, C:P, and C:Fe of heteroprokaryotes were higher than those of the phytoplankton. Limitation by oxygen followed a previous diffusion-limited model of oxygen uptake (*82*), so that aerobic heterotrophs were able to subsist down to oxygen concentrations in the nanomolar range. The oxygen demand was determined stoichiometrically following previous work (*82*, *83*).

**S2.2 Rate-affinity tradeoff and the substrate lability spectrum**

We resolved the tradeoff between the cellular optimization for maximum growth rate vs. substrate affinity with allometric scaling similar to previous work (*35*). This built upon the rate-affinity tradeoff that characterizes the trait-based resolution of diverse phytoplankton (*84, 31*). For heterotrophic prokaryotes, we considered this tradeoff to be conceptually determined predominantly by the cellular allocation of protein and energy towards one or the other optimization, rather than cell size (*21*), despite cell size also differentiating oligotrophs and copiotrophs (*18*). Maximum growth rates (at reference temperature 20°C) and half-saturation constants were parameterized as a function of cell volume $V$ as:

$$p = aV^{b}10^{c} \tag{S4}$$

for parameter $p$, where $aV^b$ encapsulated the allometric tradeoff and $10^c$ incorporated the variation in lability via the baseline maximum growth rate. In the model, $c$ varied from -1.5 to 1.5 for DOM, spaced logarithmically with five DOM pools, allowing for three orders of magnitude of variation in lability. For particulate organic matter, since it was resolved more simply, $c$ varied from -1.0 to 1.0 with three POC pools. In the illustrated simulations, we resolved two size classes of heteroprokaryotes of cell diameters 0.4 and 0.6, which were the two smaller of the three size classes resolved in previous work with one DOM pool (*35*). Including all three size classes resulted in similar biogeographies for the two larger size classes. All trait values for DOM and POM consumers are listed in Table S2.

*Implicit ecological complexity*: In the present framework, we assumed 'specialist' consumption by each functional type on one broad class of organic substrates. In previous work (*7, 80*), we demonstrated the impact of considering 'generalist' functional types able to consume more than one lability class. The solutions converged to be qualitatively the same if there was a penalty for generalist ability that scaled linearly with the number of organic pools able to be consumed. If there was a sublinear penalty or no penalty, then generalist functional types dominated, and less or no DOM accumulated. It was therefore reasonable to approximate the microbial consumption structure with these 'specialist' consuming types, since we know that recalcitrant DOM does accumulate, and that some guilds of microbes are known to specialize on recalcitrant DOM consumption. Similarly, we neglected to resolve cross-feeding among heteroprokaryotic populations. The broad organic lability classes in the model were meant to represent many (hundreds or thousands) of types of organic molecules, and the functional type consumers also were meant to represent many ecotypes of microbial consumers, including those involved in cross-feeding of organic substrates, and so the complex heterogeneity of specialist vs. generalist and primary vs. secondary consumers was considered implicitly by the model (*7, 80*).

#### S2.3 DOM production

In the model, organic matter was produced from the mortalities and excretions of all populations. Total production was partitioned into DOM vs. POM production according to a specified fraction, which differed for different functional types (e.g., 30% to DOM for the mortalities of non-photosynthetic microbial types (*85*)). POM sank at a constant rate (10 m $d^{-1}$). DOM was additionally produced from the excess hydrolysis of POM according to parameter $\alpha$ as described in (*80*) (here, $\alpha = 2$). Together with vertical mixing, the sinking and hydrolysis allowed for continued supply of both DOM and POM below the euphotic zone. Total DOM was partitioned into the multiple pools following a lognormal distribution as in previous work (*7*), meant to represent the average outcome of the multiplicative stochasticity characterizing microbial transformations of organic molecules over time and space. Hydrolysis from POM further contributed to the partitioning among DOM pools: the more labile POM pool supplied the three most labile DOM pools, the middle-range POM pool supplied the middle three DOM lability pools, and the least labile POM pool supplied the three least labile DOM pools, according to consistent weightings. Importantly, model results were qualitatively independent of these assumed distributions. A uniform distribution produced the same qualitative result with respect to the biogeographies of the microbial consumers (i.e., the fast consumers of labile DOM populate the surface and the slow consumers of recalcitrant DOM were restricted from the surface and emerged at depth due to apparent competition, along with the shift from dominance by oligotrophs to copiotrophs). However, the lognormal distribution provided reasonable quantitative relationships among the lability classes in that most of the DOM produced was of average lability, and much less was of very high or very low lability. The lognormal distribution qualitatively aligned with empirical estimates for semi-refractory, semi-labile, and labile classes (*6*; Fig. S10).

#### S2.4 Anaerobic heteroprokaryotic functional types

The model also resolved anaerobic heteroprokaryotic functional types utilizing nitrate instead of oxygen as an electron acceptor, following chemical redox-reaction-based parameterizations (*83*). Since the analysis here did not focus on the anoxic domain and anaerobic metabolisms, anaerobic heteroprokaryotic functional types were assumed to carry out full denitrification to elemental nitrogen gas, and we did not resolve the rate-affinity tradeoff among them. One anaerobic type was included as a competing consumer for each pool of organic matter (8 total: one for each of the 5 DOM pools and 3 POM pools). Trait values were set as equal to those of the copiotrophic types (though this does not matter since competition among anaerobes was not explored). Previous work (*83*) demonstrated the functional equivalence of resolving competing aerobic and anaerobic functional types compared to resolving facultatively aerobic functional types when environmental perturbations were slow relative to microbial growth rates, as was the case for the coarse-grained global model here. Anaerobic functional types (like all functional types) were introduced with equal biomasses across the global ocean domain. For the DOM consumers, anaerobic functional types were competitively excluded in the oxygenated domain due to their lower organic matter yield (*83*), including the entirely of the three illustrated transects. For the POM consumers, interestingly, the anaerobic types were not completely excluded, since the coupled supply and sinking of POM precluded complete exclusion, though they persisted at lower biomasses than their aerobic competitors. Conceptually, the persistence of these anaerobic heterotrophs in the oxygenated ocean may meaningfully represent those inhabiting the anoxic niches inside organic particles. Because the anaerobic POM consumers persisted, we considered the sum of aerobic and anaerobic POM-consumer biomasses in the analysis.

#### S2.5 Impact of temperature

All metabolic rates (including growth, grazing, and mortality), were modified as a function of temperature (non-dimensional $\gamma_T$) using a formulation that followed the Arrhenius equation (*86*):

$$\gamma_T = \tau \exp\left(A_E\left(\frac{1}{T} - \frac{1}{T_0}\right)\right) \qquad \text{(S5)}$$

where $T$ is the ambient temperature (K), $T_0$ is a reference temperature ($T_0$ = 293.15 K), $A_E$ regulates the temperature modification ($A_E$ = −4000 K), and τ normalizes the maximum value (τ = 0.8; unitless). This formulation allowed for direct comparison with the genome-based maximum growth rate estimates when setting the optimal growth temperature to 20°C. Following this common approach, we did not model functional types with different temperature optimums. Rather, the heteroprokaryotic functional types were considered as aggregates of many clades with different temperature optimums. Variation in temperature optimum may indeed explain much of the observed diversity, particularly among ecotypes within clades. Future work could explore the finer-scale diversity resulting from optimization to different water temperature ranges.

#### S2.6 Grazing structure

We incorporated the DOM-consuming heteroprokaryotic functional types framework into the size-based grazing framework employed by the Darwin-MITgcm ecosystem model (*34*). For POM consumers, grazing and other mortality dynamics on particles are uncertain, but grazers are likely distinct from those of the free-living populations, and so grazing of POM consumers was resolved implicitly using a higher density-dependent (quadratic mortality) coefficient. As in previous work (*35*), the smaller oligotrophic functional types had their own separate grazer, while the larger copiotrophic functional types shared a grazer with similarly sized picophytoplankton. Critically, our results did not depend on the size-based particulars of the grazing scheme, but only on the presence of a shared grazer among the microbial populations (see similar results in the model sensitivity experiment with one grazer in Fig. S19). As long as the slow-growing consumers of recalcitrant DOM shared a grazer with faster-growing microbial populations at the surface, and grazing pressure was sufficient such that the losses to grazing were higher than the maximum growth rates of the populations, the slow-growing consumers were excluded from the surface.

### S3 Multivariate and other statistical analyses

Here we describe the analysis of Bray-Curtis dissimilarity matrices of both sequencing data and modeling output using non-metric multidimensional scaling (NMDS), hierarchical clustering, and several other nonparametric multivariate statistics. All analyses were performed using R version

4.1.3, except for the rank correlation analysis, which used Python's SciPy library (version 1.5.2). For the P16 N/S metabarcoding dataset, the analysis focused on the 21 taxonomic guilds. Relative abundances were re-calculated to generate a new compositional matrix consisting of the 21 guilds for each sample along the P16 transects (n = 189). This was used to calculate a pairwise Bray-Curtis dissimilarity distance using the 'vegdist (method = "bray")' function.

NMDS ordination (function: 'metaMDS (distance = "bray", autotransform = F, wascores = T)' was performed to visualize the pairwise Bray-Curtis dissimilarities. The goodness of fit statistic (function 'goodness') was evaluated with Shepard diagrams (function 'stressplot'). The NMDS stress, with large values (e.g., > 0.2) indicating poor fits, was specified for each ordination plot in Fig. 3 and Fig. S16. We also used the function 'envfit (permutations = 999)' to fit the 21 functional guilds as environmental variables on the ordination space. These vectors indicated the direction of maximum change in the ordination space, with vector length scaled by the strength of the correlation between the environmental variable and ordination scores. We also specified, using different line styles, the classification of each of the guilds according to the copiotrophy index, conducted independently from this analysis (i.e., our results from the genome-based analysis as depicted in Fig. 4). All plots were generated using the 'ggplot2' package. The functions 'vegdist', 'metaMDS', 'goodness', 'stressplot', and 'envfit' were from the 'vegan' package.

Additionally, we grouped samples by depth bins (k = 2, 3), biogeographic regions (k = 5), Longhurst provinces (k = 10), and a parallel output of an agglomerative hierarchical clustering approach with corresponding number of clusters (e.g., k = 2, 3, 5, 10). For clustering, we used the 'hclust (method = "ward.D2")' function from the 'stats' package to carry out the Ward's minimum variance. We also used the 'fviz nbclust (method = "wss")' function from the 'factoextra' package to help visualize and estimate the optimal number of clusters using the total within sum of square method. The analysis of similarities (ANOSIM), function 'anosim' from the 'vegan' package, was performed to test for a significant statistical difference between the two or more groups of clusters devised using oceanographic context (e.g., depth bins, biogeographic regions) or derived from hierarchical clustering algorithms. We also carried out the permutational multivariate analysis of variance (PERMANOVA), function 'adonis' from the 'vegan' package, to provide a percent estimate for how much of the variation of the community composition (e.g., Bray-Curtis distances) was explained by the groups or clusters.

*Application to modeled communities:* We reproduced the above workflow with the model simulation by subsampling along a parallel Pacific Ocean transect at colocalized geographical coordinates and depth bins in which empirical observations were made, resulting in a comparable number of model data points (n = 174). As in the data, we analyzed just the heteroprokaryotic functional types. We excluded the types below a biomass threshold of $10^{-6}$ mmol C $m^{-3}$ (i.e., functionally extinct). To reduce influence from distinctions among the three particle-associated (POM-consuming) types, since we did not focus on understanding this diversity, we aggregated these three into a single POM-consuming class. Also, as in our analysis of biogeographies (Fig. 2, Fig. S12), we aggregated the three labile-DOM-consuming oligotrophs and the three labile-DOM-consuming copiotrophs. (When we did keep all of these types distinct in the analysis, the NMDS results did not change significantly, but these aggregations allowed for a simpler and clearer interpretation and presentation.) In sum, we analyzed the communities with seven modeled types (Fig. S12): a POM-consuming type, a labile-DOM-consuming copiotroph and oligotroph, a semi-labile-DOM-consuming copiotroph and oligotroph, and a semi-refractory-DOM-consuming copiotroph and oligotroph. The biomass values for these model community members were converted into a compositional data matrix with % relative abundance values, and the Bray-Curtis dissimilarity matrix was calculated from the model simulation.

The resulting Bray-Curtis distances were visualized on a NMDS plot, with parallel supplemental analysis using the 'envift', 'anosim', and 'adonis' functions, to objectively investigate which model functional types are driving community composition and how similar groups or clusters (e.g., depth, biogeographic regions, etc.) might play a role in structuring the emergent heteroprokaryotic community in the model seascape (Fig. S16). The sample size along the model P16 N/S transects was resampled with some redundancy ($n$ = 189) to match the dimensions of the empirical observations for further analyses. This included carrying out the Mantel test, function 'mantel' (method = "spearman") from the 'vegan' package, to test the similarity between Bray-Curtis indices of parallel samples in the molecular dataset and model simulation. Furthermore, we used the 'vegan' package to carry out a Procrustes analysis, function 'procrustes' (symmetric = T), and subsequent correlation in a symmetric Procrustes rotation, function 'protest' (score = 'sites', permutations = 999), to test the correlation in patterns of community structure between the molecular and model seascape using co-localized samples from the respective NMDS ordination plots. Fig. S17 illustrates the residuals of the rotation as a function of geography.

*Rank correlation analysis*: We compared the ordering of the relative abundances of comparable groups for the modeled and empirical communities. Because comprehensive and comparable groups were essential for this analysis, we were limited to analyzing four groups (illustrated in Fig. S14): (i) All "Surface, low-latitude/ubiquitous" guilds compared to the modeled labile-DOM-consuming oligotroph, (ii) All "surface, high-latitude" guilds, plus *Enterobacterales,* (because our analysis showed this guild as an extreme, fast-growing copiotroph) compared to the modeled labile-DOM-consuming copiotroph, (iii) All "deep, ubiquitous" guilds compared to all modeled recalcitrant-DOM-consumers, and (iv) All "particle-like" guilds compared to modeled POM consumers. Because NMDS of the empirical communities showed an optimum of three clusters, we also conducted the rank correlation analysis when combining groups iii and iv into one deep-peaking group to give a total of three groups. We computed the Spearman's rank correlation coefficient for each sampling location (using the 'spearmanr' function from Python's SciPy library, version 1.5.2) to assess the degree to which the ordering of the relative abundances of these groups was similar between data and model (larger colored dots in Fig. S18). A coefficient of 1.0 meant that the ordering of the groups, in terms of relative abundance, was the same for both data and model even though the magnitudes of the relative abundances differed. We also assessed whether the model matched the observed dominant group, i.e., whether the same group was ranked as first (smaller black dots in Fig. S18).

### Supplementary Text

**Robustness of model results** The heteroprokaryotes consuming recalcitrant DOM were partially or completely excluded from the surface layer across the plausible parameter space. A functional type was excluded because its population (biomass) loss rate exceeded its maximum growth rate. Thus, the model results depended on the assumption that the consumers of recalcitrant DOM grow more slowly than consumers of labile DOM. Partial exclusion was possible because vertical mixing and other physical transport supplied low rates of biomass to areas in which they weren't able to survive in isolation. The only cases in the model in which recalcitrant DOM consumers subsisted sustainably in the surface were when the model resolved very low rates of primary productivity in the center of oligotrophic gyres. When this happened (when productivity rates were much lower than observed rates in the gyres) surface grazing rates were low enough for the semi-labile consumers to subsist (for example, see Fig. S20). Surface exclusion of the slow growers also still occured when there was no assumed difference in cell size between copiotrophs and oligotrophs, such that one zooplankton grazed upon all DOM-consuming heteroprokaryotic functional types (Fig. S19). However, surface exclusion in this latter model configuration (with one grazer of all DOM-consuming heteroprokaryotes) held only as long as the copiotrophs consuming recalcitrant DOM grew more slowly than the oligotrophs consuming labile DOM. If, alternatively, parameter values were such that copiotrophic growth on recalcitrant DOM was faster than oligotrophic growth on labile DOM, results would hold only when the larger copiotrophs and smaller oligotrophs retained their distinct grazers, and thus when the assumed difference in cell size between oligotrophs and copiotrophs was retained (as in the default model).

**Model experiments** Figs. S19–S22 illustrate results from three different model sensitivity experiments. First, Fig. S19 illustrates results in which all heteroprokaryotic consumers (oligotrophs and copiotrophs) were grazed upon by one shared grazer, rather than assuming that smaller oligotrophs have a different grazer. This showed that results held regardless of the assumptions about heteroprokaryotic cell size and its relationship with grazing. As long as the slow-growing heteroprokaryotes consuming RDOM shared a grazer with a sufficiently abundant, faster-growing population, they were excluded from the surface. Second, Fig. S20 illustrates results when there were no explicit grazers at all for heteroprokaryotes. For this experiment, the quadratic mortality constant was increased to account for grazing implicitly, such that growth and remineralization rates remained similar. (Results would be qualitatively the same if each functional type was consumed by its own individual grazer.) This experiment resulted in no surface exclusion of any functional types, illustrating clearly that shared grazing was the mechanism for the exclusion. Third, illustrated in Figs. S21–S22, we determined why the model captured the greater ubiquity of the SAR324 model analog (the semi-labile-consuming copiotroph) relative to the SAR202 analog (the semi-refractory-consuming copiotroph). To do so, we conducted a model experiment in which the supply rate of semi-labile and semi-recalcitrant DOM was equal, rather than following the default log-normal distribution in Fig. 1. With this equal supply, the only difference between the consumers was the difference in lability that corresponded to the baseline maximum growth rate. Results showed the same biogeographical pattern as in the default model, confirming that it was the difference in maximum growth rate that set the different biogeographies. Because it was complicated to assure equal supply of the two DOM classes through the fluxes of excess hydrolyzed POM, we set up a model configuration with no excess hydrolysis ($\alpha = 1$; Fig. S21) for direct comparison with the experiment in Fig. S22.

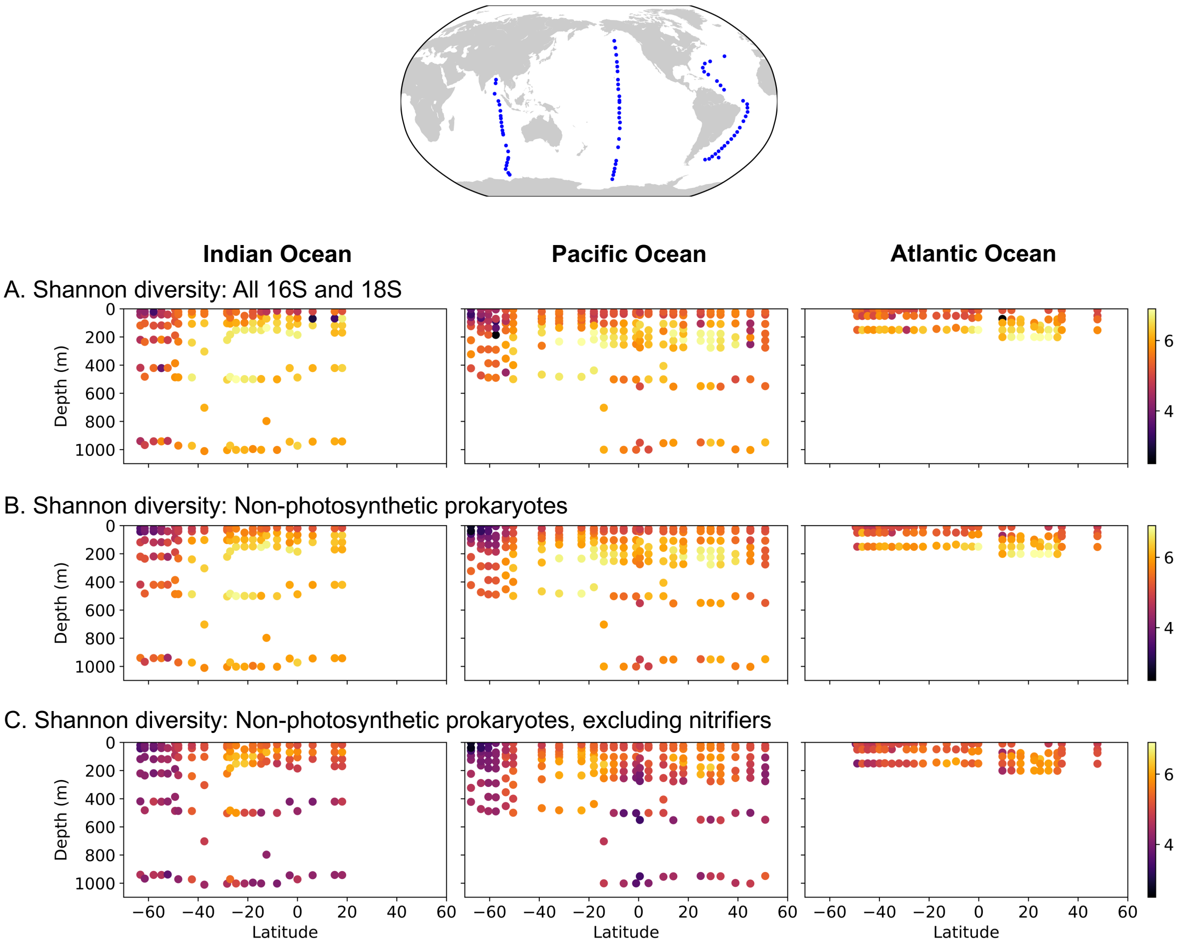


**Figure S1:** Diversity patterns along the three ocean transects, showing the ubiquitous increase in diversity from surface waters (roughly 0-150 m) to deeper waters (peaking at roughly 200-500 m). (A) The Shannon diversity index of the whole community according to 16S and 18S metabarcoding data. (B) The Shannon diversity index of the non-photosynthetic prokaryotic community (including nitrifying microorganisms) according to 16S metabarcoding data. (C) The Shannon diversity index of just the heteroprokaryotes community (excluding nitrifying microorganisms). Nitrifying organisms are identified using the keywords “Nitroso” and “Nitrosp”.

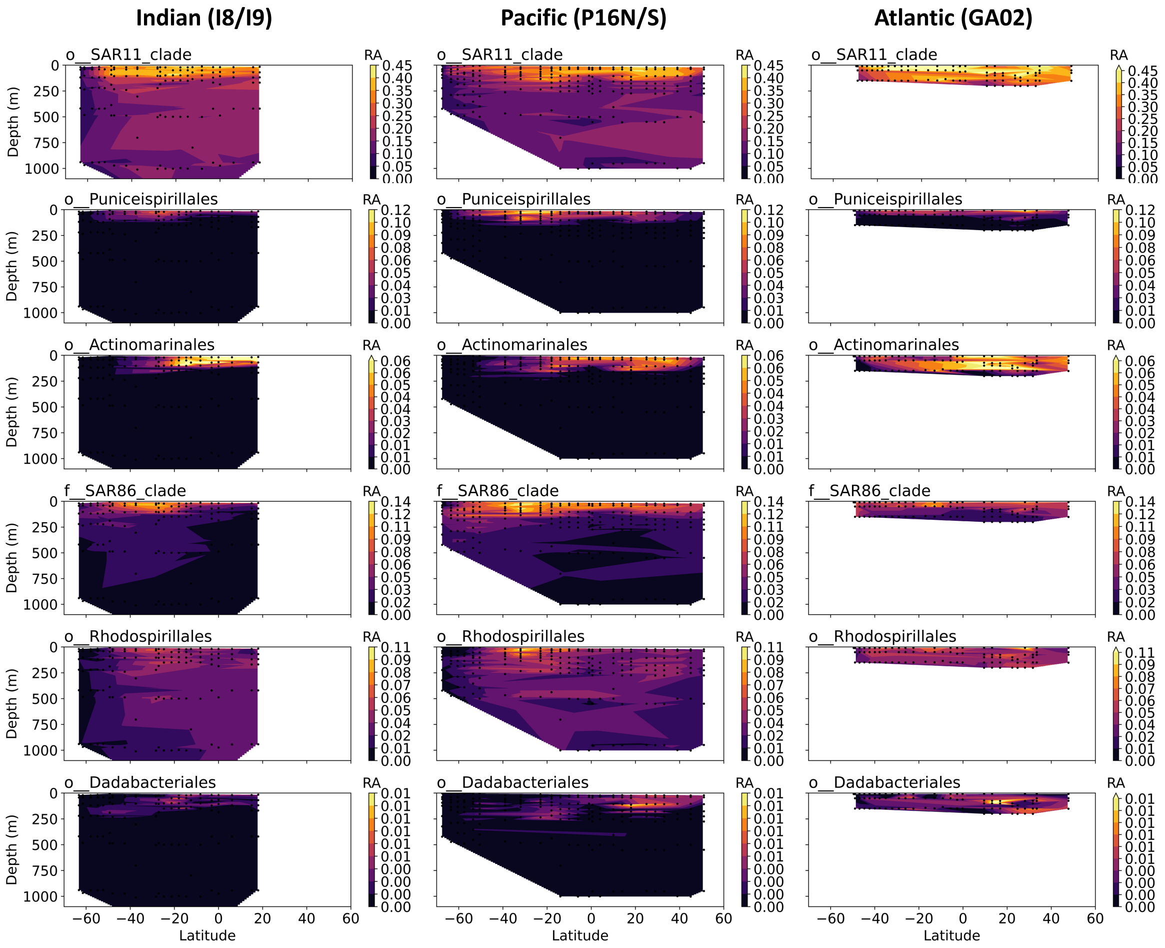


**Figure S2:** The relative abundances (RA) of the six *surface, low latitude/ubiquitous* heteroprokaryotic taxonomic guilds with peak relative abundances across the surface, ubiquitously, and predominantly at lower latitudes (roughly 40$^o$N to 40$^o$S).

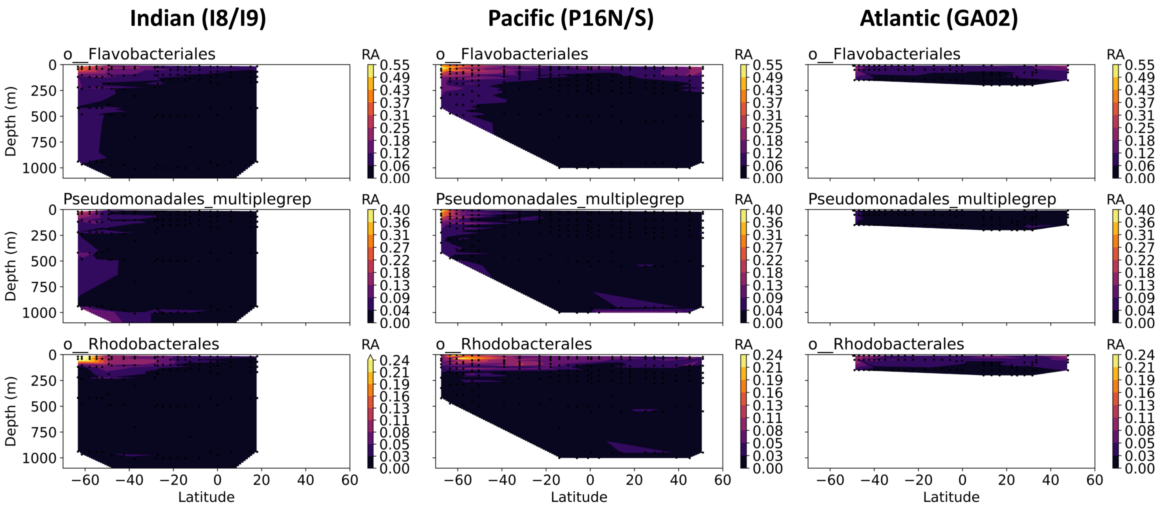


**Figure S3:** The relative abundances (RA) of the three *surface, high latitude* heteroprokaryotic taxonomic guilds with peak relative abundances at high latitudes. Note that the Southern Ocean was sampled during the productive summer season, unlike the higher northern latitudes.

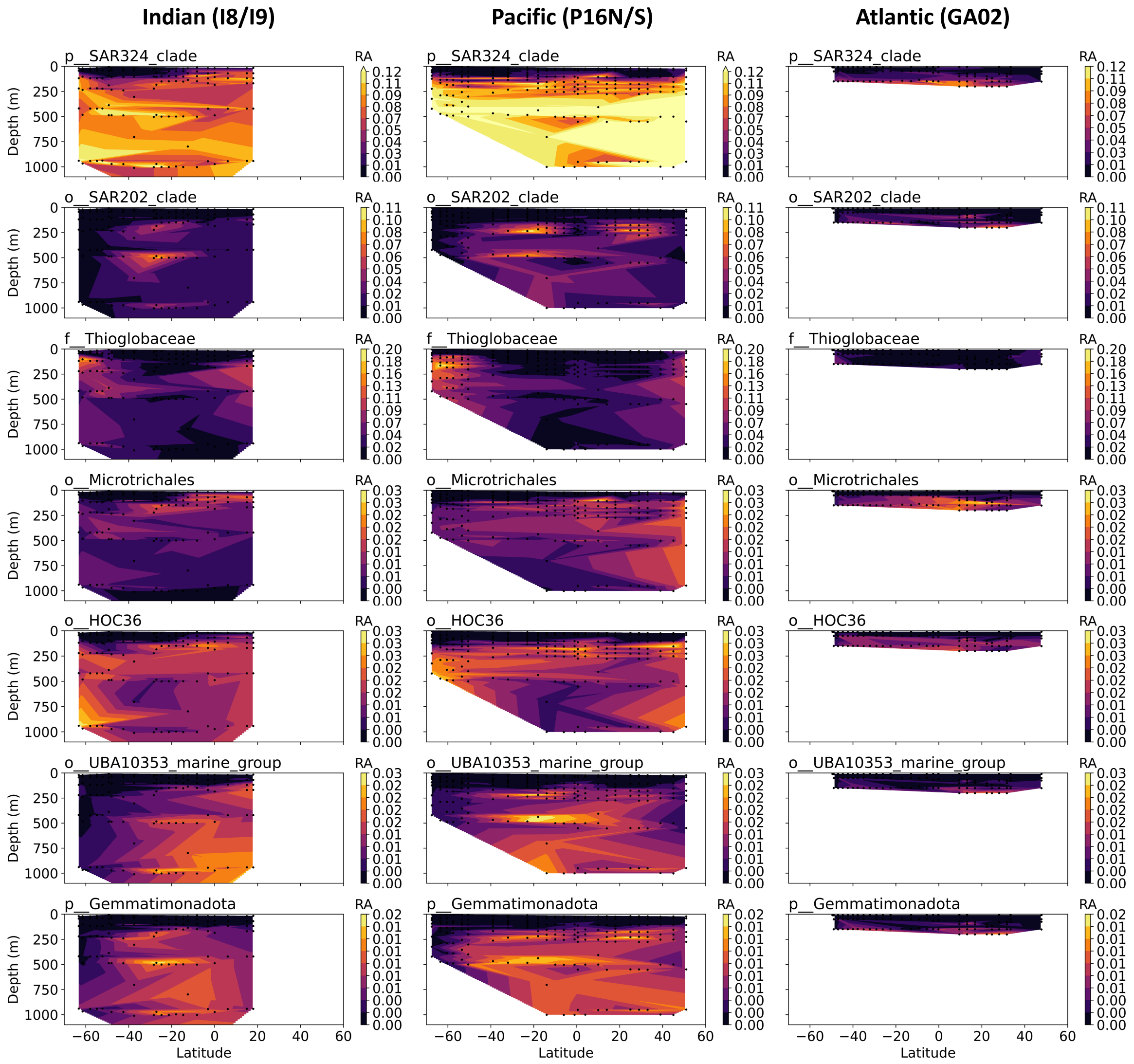


**Figure S4:** The relative abundances (RA) of the seven *deep, ubiquitous* heteroprokaryotic taxonomic guilds with peak relative abundances in the deep ocean with little abundance in the surface.

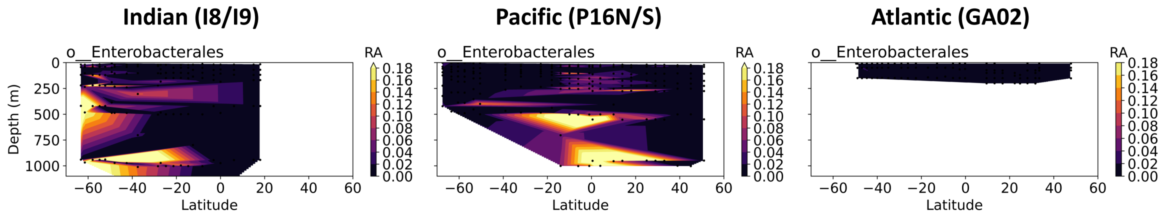


**Figure S5:** The relative abundances (RA) of the one *deep, sporadic* heteroprokaryotic taxonomic guild with peak relative abundance that appears sporadically in the deep ocean (i.e. very high RA in some areas, and very low RA in others).

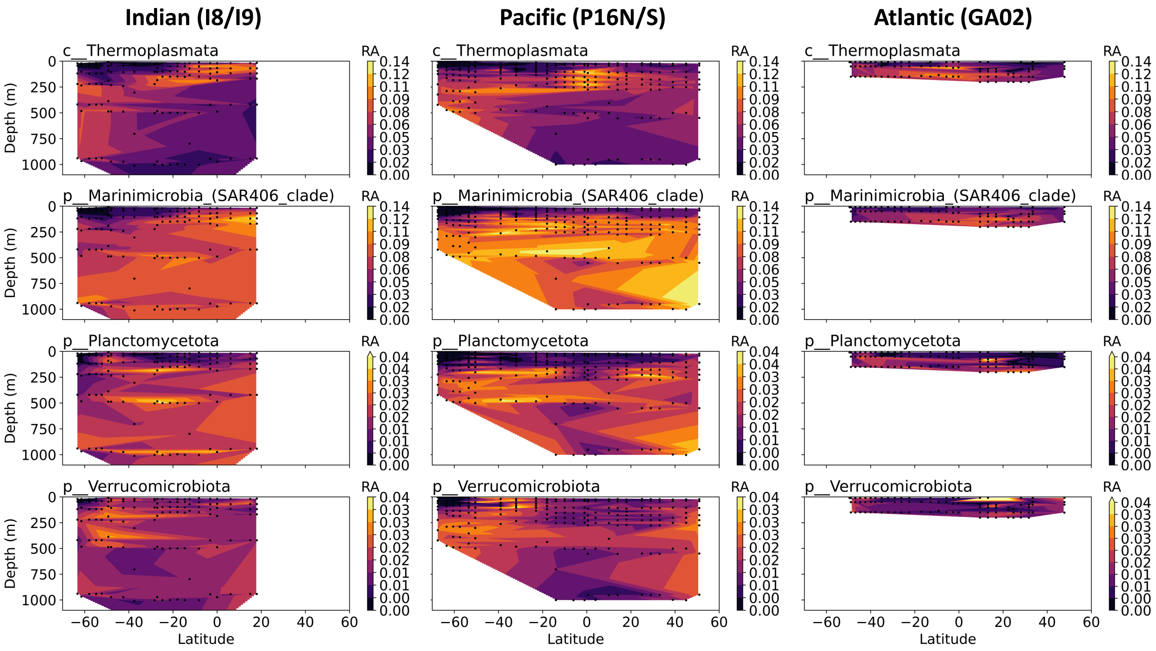


**Figure S6:** The relative abundances (RA) of the four *particle-like* heteroprokaryotic taxonomic guilds with relatively ubiquitous relative abundances, peaking at depth in general but also with more substantial relative abundances in the surface than those in Fig. S4. These distributions match most closely with the modeled consumers of particulate organic matter (see Fig. S13).

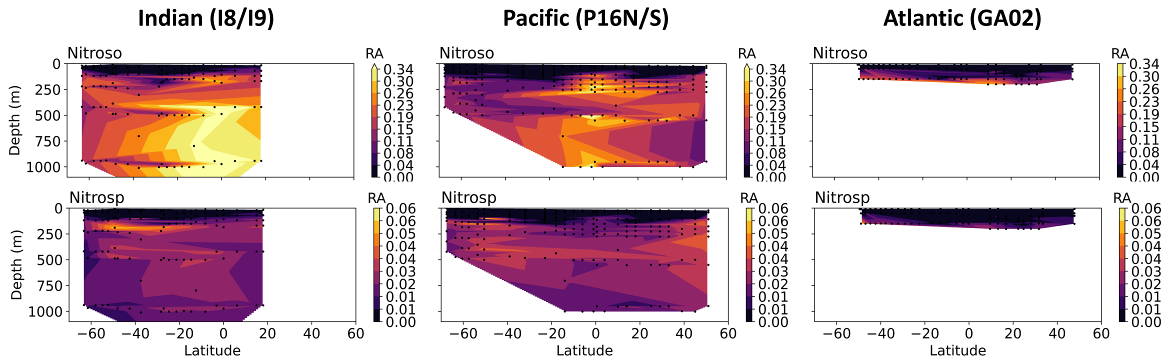


**Figure S7:** The relative abundances (RA) of the ammonia-oxidizing (keyword “Nitroso”) and nitrite-oxidizing (keyword “Nitrosp”) guilds.

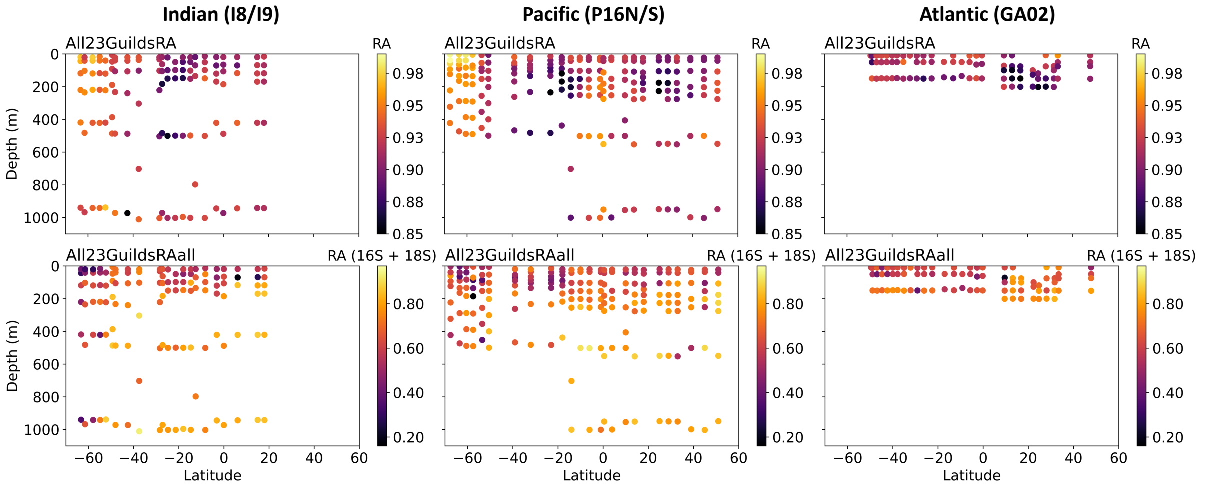


**Figure S8:** The fraction of the non-photosynthetic prokaryotic community covered by the 21 guilds plus the two nitrifier guilds. Top row: Fraction with respect to the non-photosynthetic prokaryotic subset of the community. Bottom row: Fraction with respect to the full community (all 16S and 18S). With respect to the non-photosynthetic prokaryotic subset, the average coverage is 92% for the Indian and Pacific transects and 91% for the Atlantic transect.

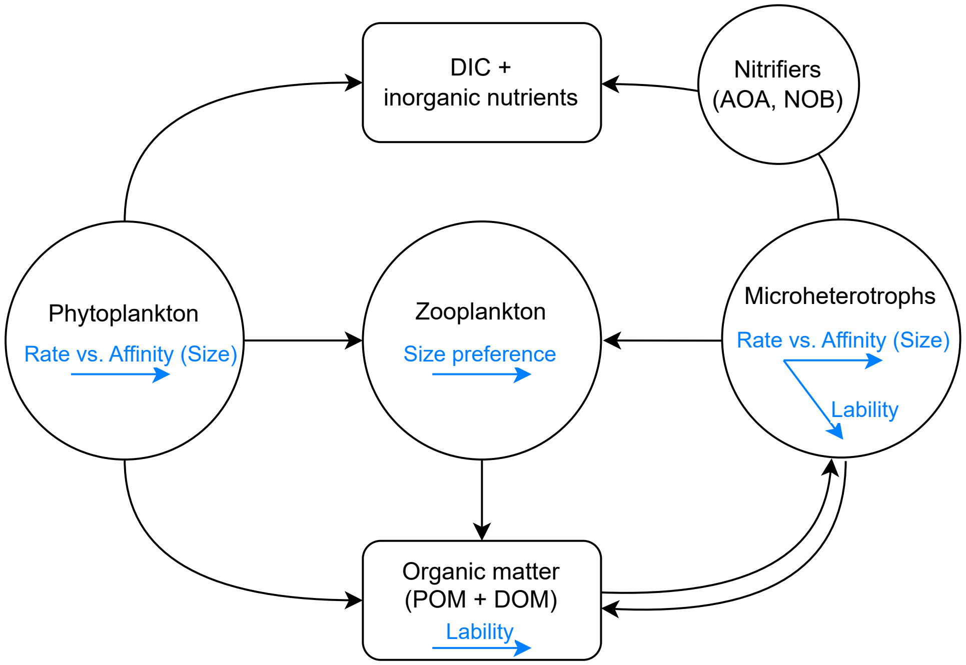


**Figure S9:** Schematic of the configuration of the marine ecosystem model (Darwin-MITgcm).

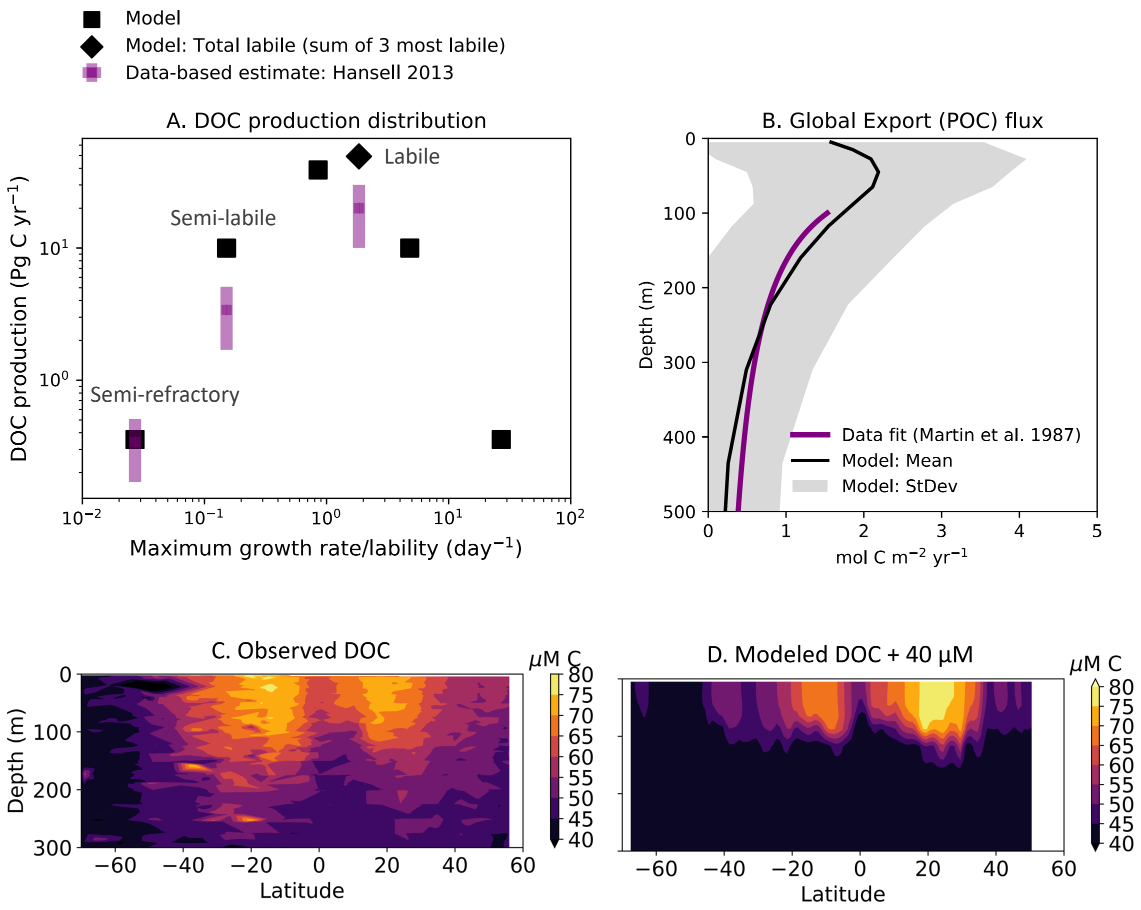


**Figure S10:** Modeled organic carbon dynamics compared to observations. (A) Modeled dissolved organic carbon (DOC) production (global annual average) compared to empirical estimates of labile, semi-labile, and semi-refractory classes in Hansell 2013 (*6*), which shows that our theoretical resolution of DOC production (i.e., the lognormal distribution across the lability spectrum) qualitatively captures the empirically inferred distribution. Total modeled labile DOC production is the sum of the three classes that are functionally labile in the surface (*7*) (see Fig. S15), and is plotted against the production-weighted lability of those three classes. (B) Modeled annually and globally averaged POC flux (with mean and standard deviation weighted by surface area) compared to empirical estimates in Martin et al. 1987 (*90*), demonstrating that the model qualitatively and somewhat quantitatively captures not only the magnitude of the global export flux (10 Pg C yr$^{-1}$ at 120 m depth; Fig. S11), but also its vertical structure. (C and D): Observed and modeled DOC concentrations along the Pacific transect (P16N/S). The model does not resolve the large refractory DOC pool, and so 40 $\mu$M is added to the modeled total just in this illustration to quantitatively compare with observations.

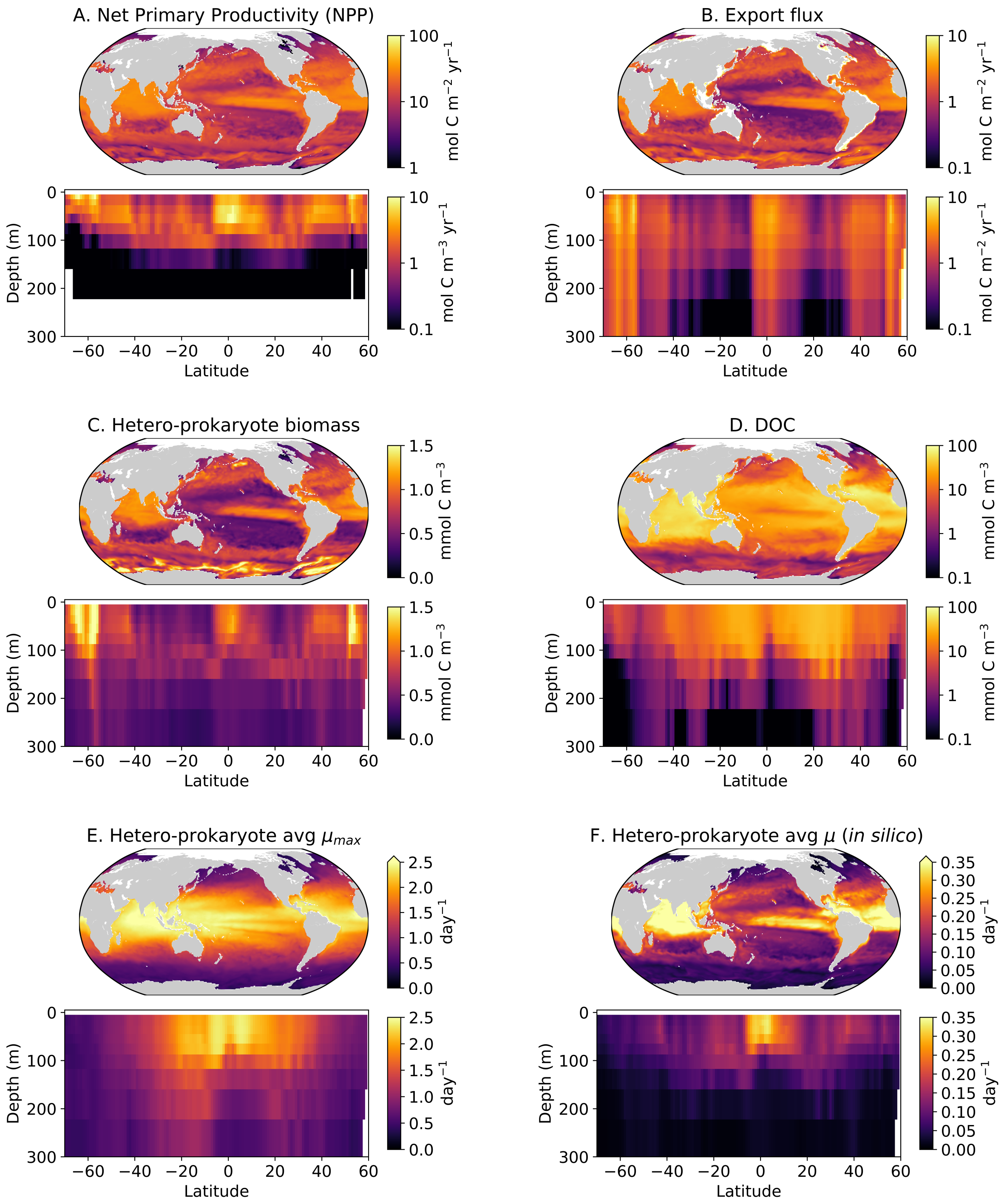


**Figure S11:** Annually averaged global model solutions, illustrating lat-lon relationships and along the P16 transect. (A) net primary production (NPP; vertically integrated for the lat-lon plot), (B) the export flux of sinking organic matter (at 100 m for the lat-lon plot), (C) total heteroprokaryotic biomass (surface concentration for the lat-lon plot), (D) total dissolved organic carbon (DOC; surface concentration for the lat-lon plot), (E) heteroprokaryotic community-averaged (abundance-weighted) maximum growth rate ($\mu_{max}$), accounting for modulation by temperature (Eqn. S5), and (F) heteroprokaryotic community-averaged (abundance-weighted) *in silico* growth rate ($\mu$). In surface waters, modeled growth rates are on average of order 0.1–1 day$^{-1}$, which is consistent with a comprehensive analysis of observations (*91*). Modeled global integrals: NPP is 63 Pg C yr$^{-1}$, similar to other global and remote-sensing based estimates (*3*), and the export flux at 120 m depth is 9.7 Pg C yr$^{-1}$, in the range of other estimates (5–11 Pg C yr$^{-1}$) (*92–94*). Bacterial biomass concentrations are roughly similar in magnitude to those of phytoplankton, consistent with observations (*95*).

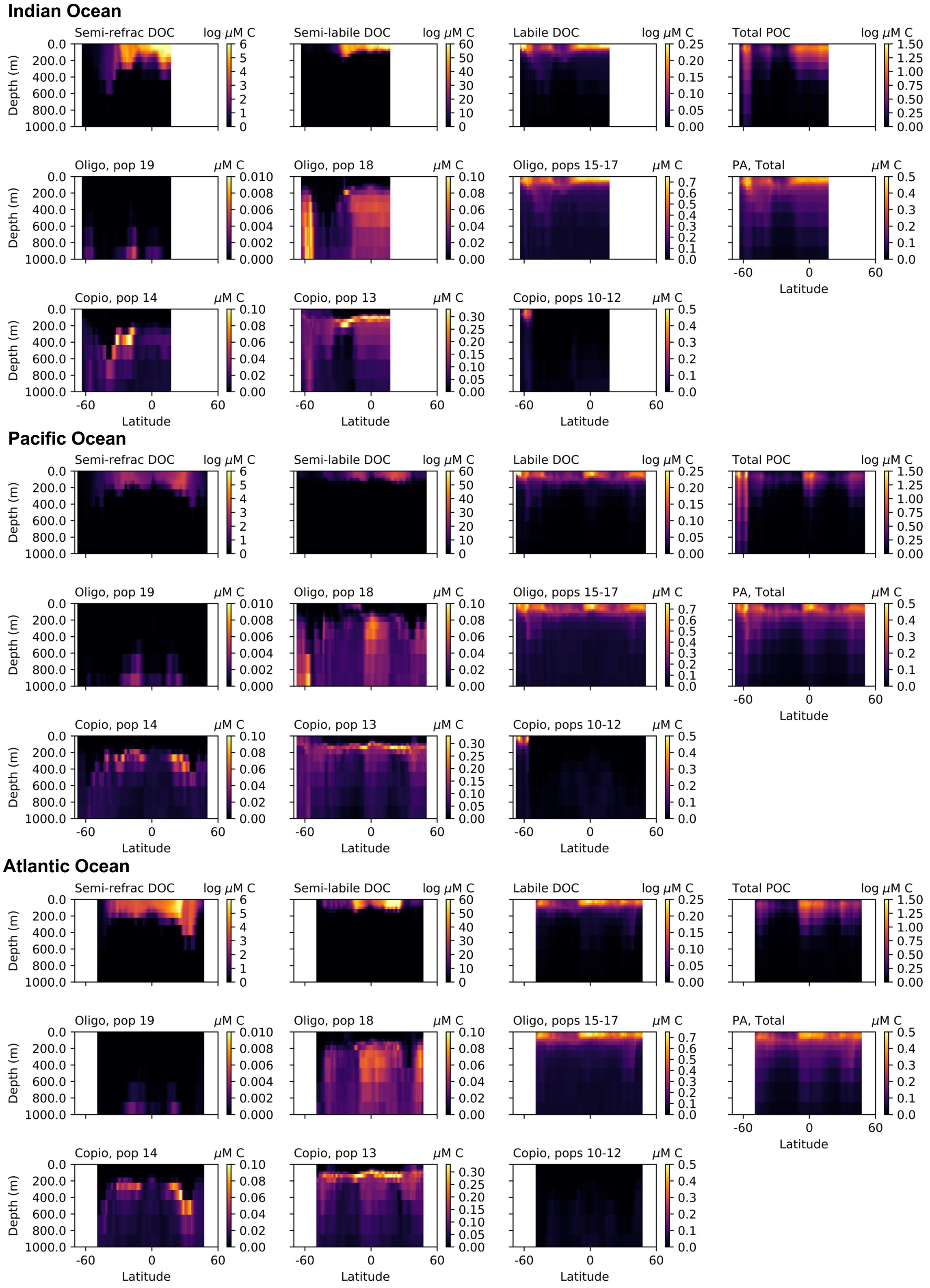


**Figure S12:** Modeled DOC and heteroprokaryotic consumer biomass across the three transects. For each of the DOC classes, the biomasses of the oligotroph and copiotroph competitive consumers are plotted underneath. For total POC (sum of 3 classes), the biomasses of all particle-associated (PA) types are plotted underneath.

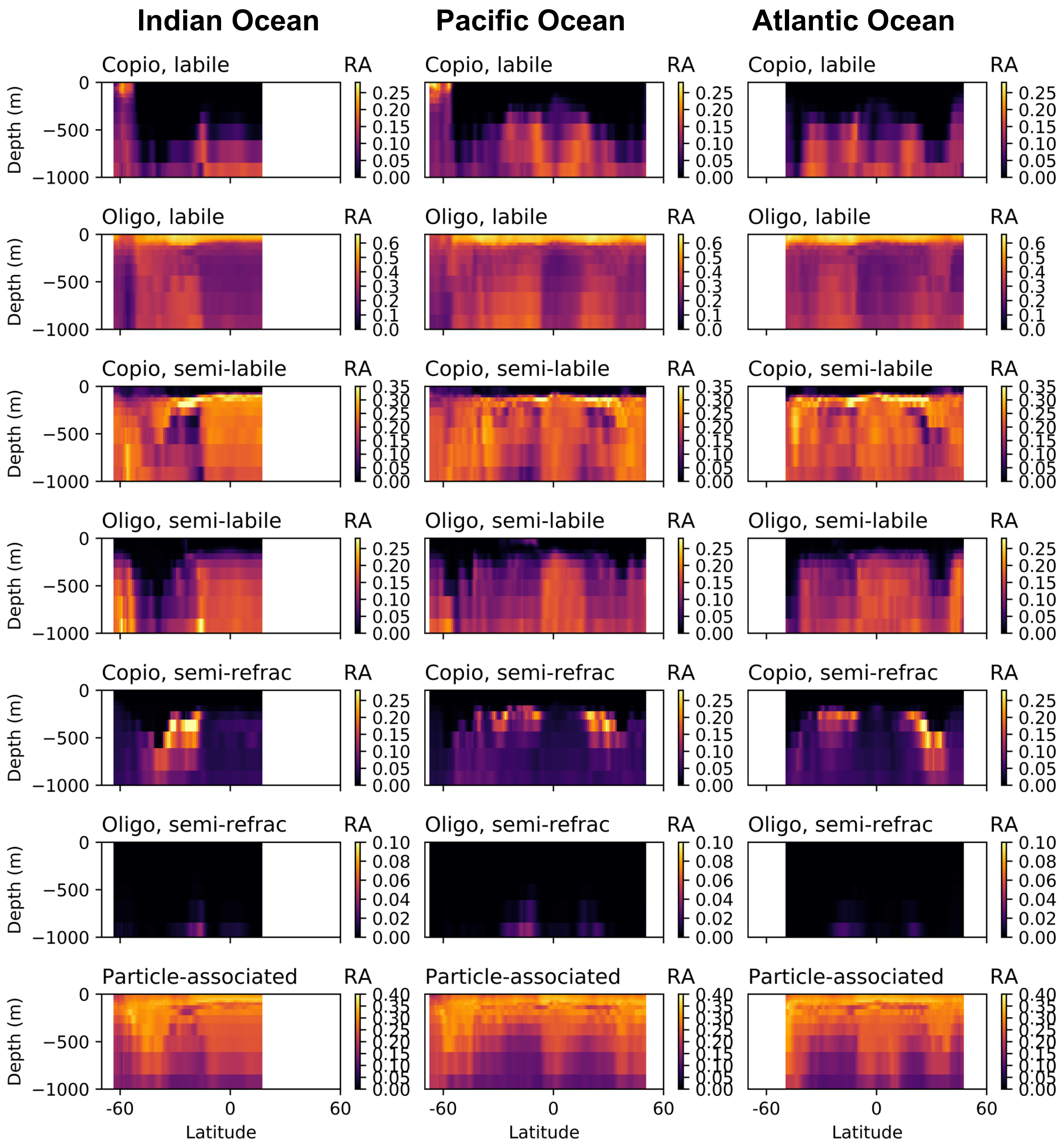


**Figure S13:** Modeled relative abundances (RA) of heteroprokaryotic consumer biomass along the three transects, labeled by their life strategy ("Copio" for copiotrophic consumers of DOM, "Oligo" for oligotrophic consumers of DOM, or Particle-associated consumers of POM) and by the type of organic matter that each consumes in the model (labile DOM, semi-labile DOM, or semi-refractory DOM, or POM). At very deep depths (below 300 m), total biomass becomes very low, and so the RA of some types (the labile DOM-consuming copiotroph, for example) becomes higher again despite much lower biomass, because fitness differences become less significant as biomass turnover rates become very small.

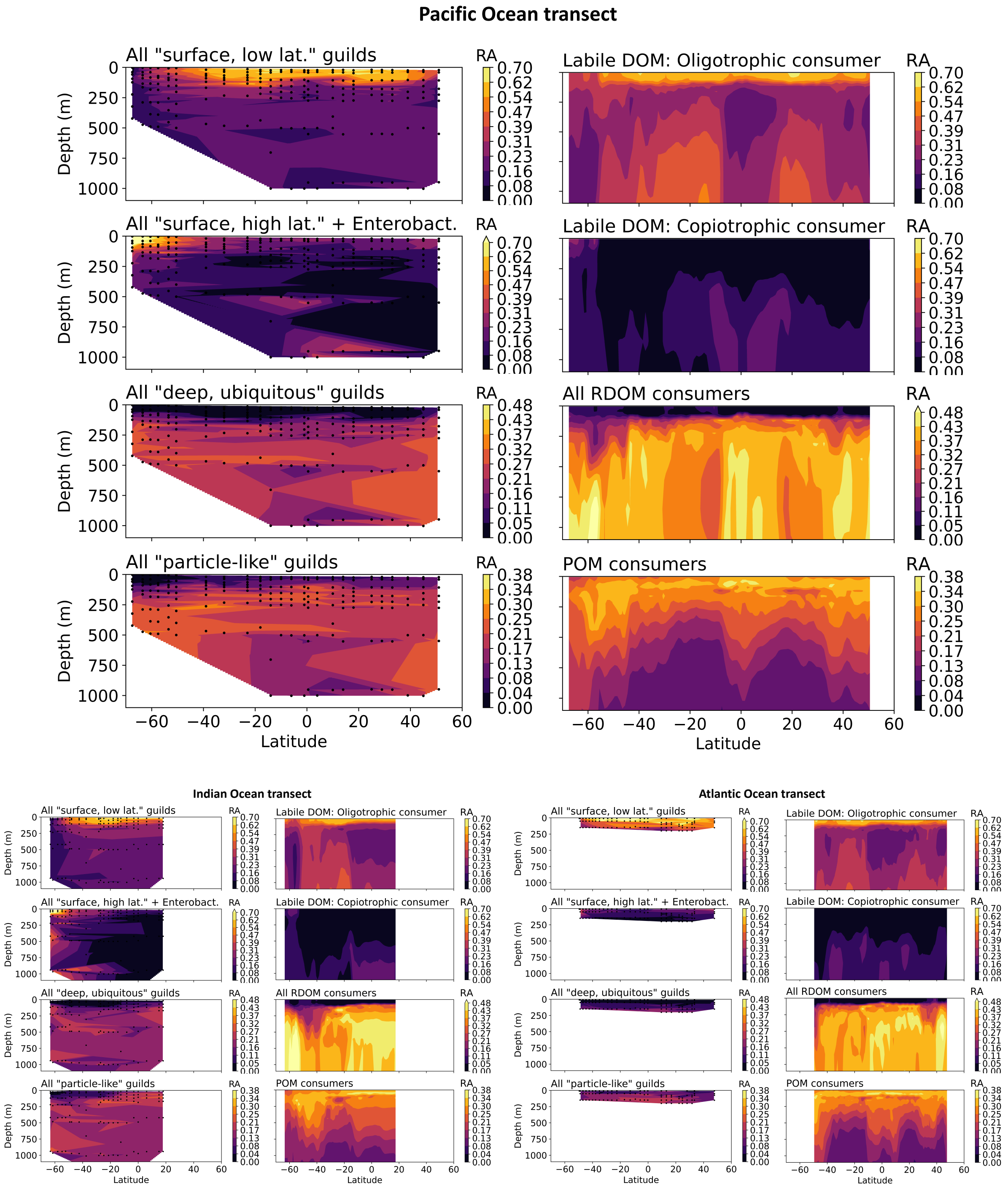


**Figure S14:** Summed relative abundance (RA) of four comparable groups of guilds and functional types. The Pacific transect is emphasized because that was used for the rank correlation analysis (Materials and Methods S3). Here, we include the deep-peaking, copiotrophic guild *Enterobacterales* along with the *surface, high latitude* guilds because we find that its biogeography and genome-inferred function (the most extreme copiotroph value with highest maximum growth rate) is consistent with the trait-based description and model-emergent biogeography of the modeled labile-DOM-consuming copiotroph.

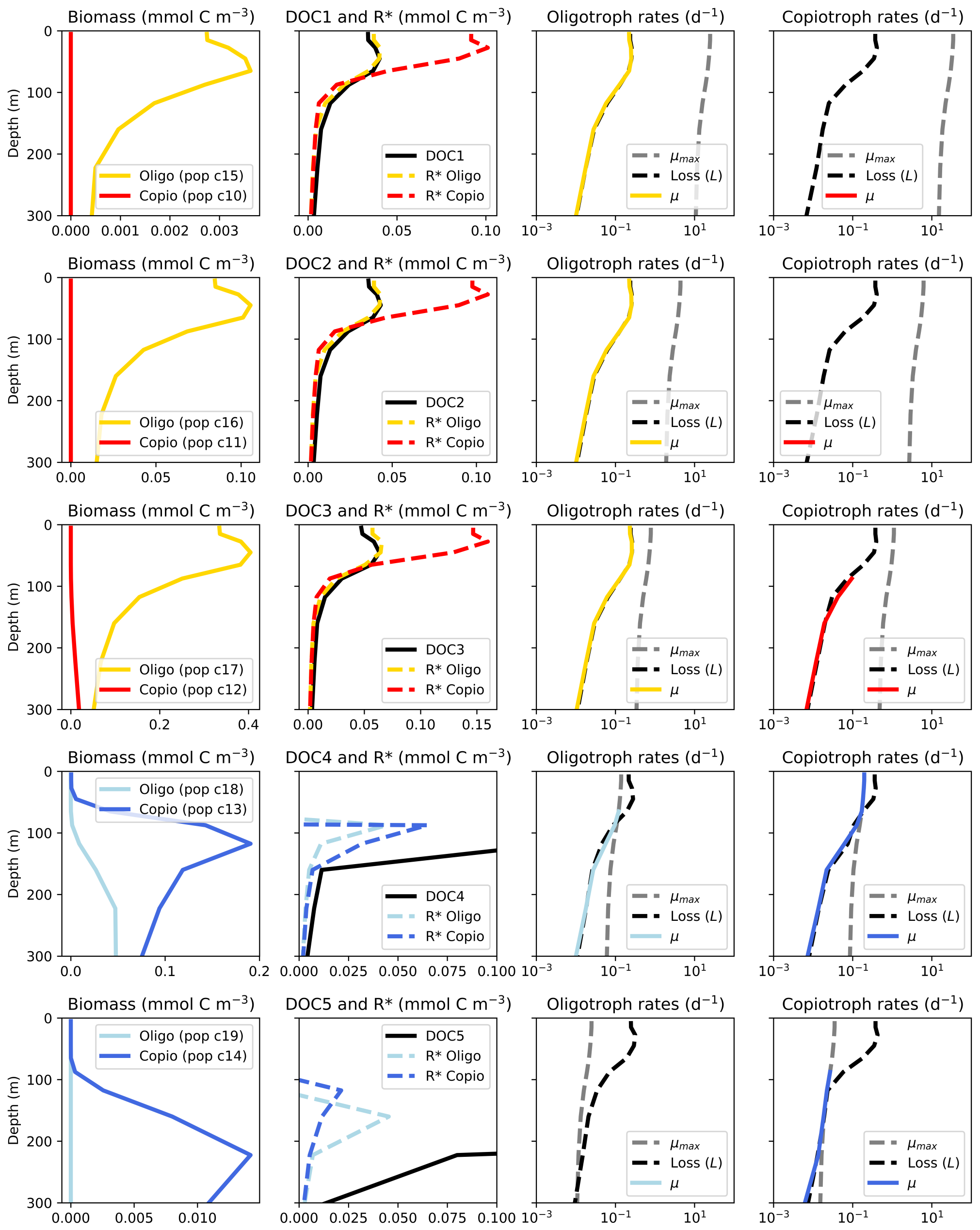


**Figure S15:** Model solutions and analysis of the ecological dynamics of DOC and heteroprokaryotic consumers. Illustrated solutions are for one representative water column location, at 10$^o$N along the modeled Pacific transect. Each row illustrates the dynamics associated with each of five DOC classes (labeled in Column 2). Column 1: The resulting biomasses of the competing oligotrophic and copiotrophic functional types. Column 2: The concentration of the DOC class and the resource subsistence concentrations (R* concentrations) for each of the two consumers. Columns 3 and 4: The maximum growth rate ($\mu_{max}$), specific loss rate, and the resulting steady-state *in silico* growth rate ($\mu$) of the competing oligotroph and copiotroph functional type, respectively. While differences in $\mu_{max}$ vs. affinity determine the fitness difference, and thus the competitive outcome, of the competing pair, the resulting *in silico* $\mu$ is similar because $\mu$ is set by the loss rates in the ecosystem. The parameters setting these loss rates (mortality and grazing constants) are extremely uncertain, and so in order to not introduce additional (uncertain) differences between functional types, we set these loss parameters to be the same for all types. *Functionally labile aggregation*: DOC class 1, 2, and 3 are dynamically classified as functionally labile because their concentrations match the R* concentrations (*7*), and so in all other model plots these types are aggregated into one labile DOC class. Biomasses of the consumers are also aggregated in the other plots (into one labile-DOM-consuming oligotroph and one labile-DOM-consuming copiotroph), because their relative distributions are nearly identical. The absolute concentrations of the consumers differ because of the difference in the production rate among the three labile classes.

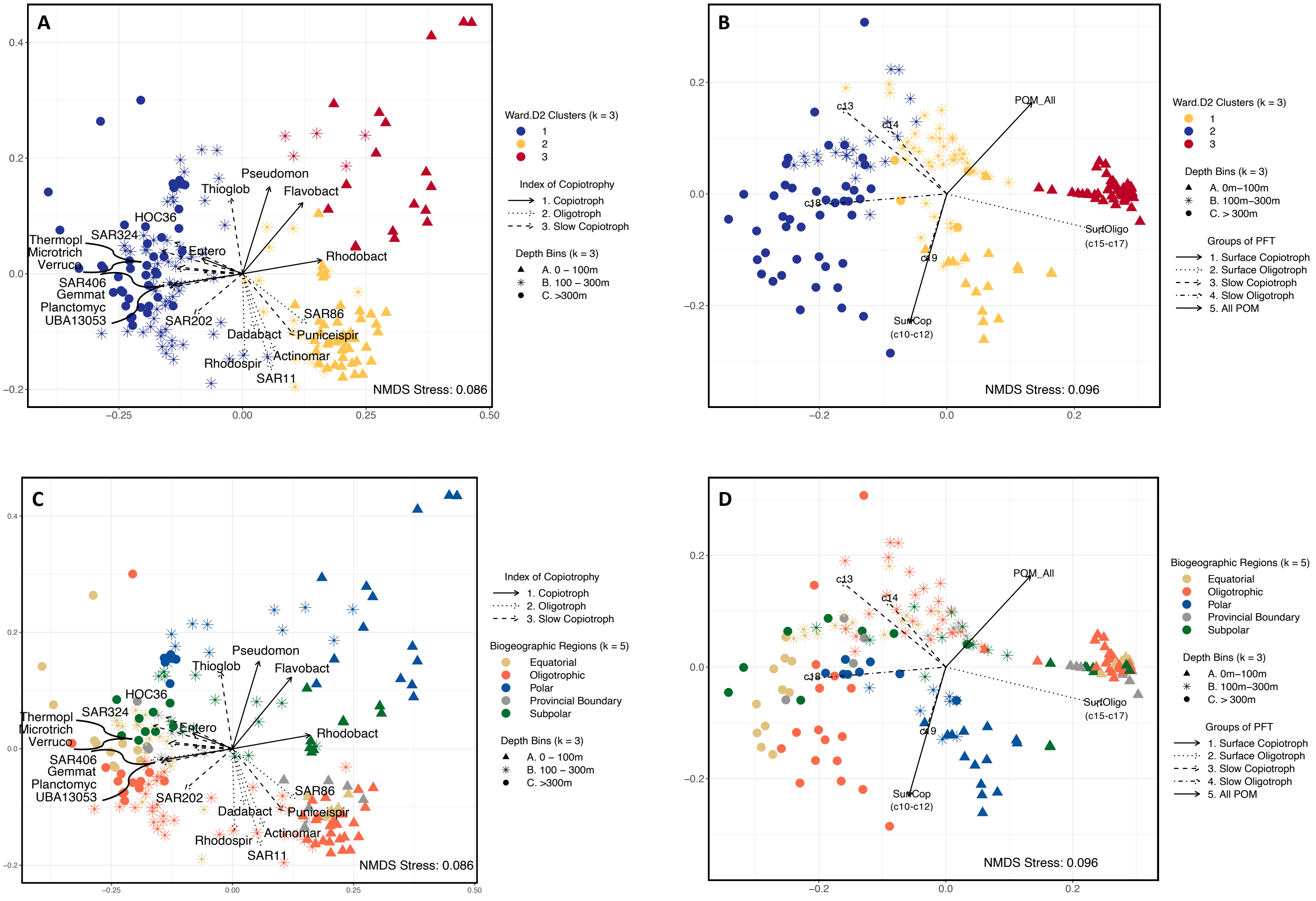


**Figure S16:** Results of the non-metric multidimensional scaling analysis (NMDS) for the empirical and modeled heteroprokaryotic communities along the P16 transect. The axes are determined from analysis of the co-located sampling and modeled locations (i.e. the 189 locations), and the arrows representing each guild or model functional type are then plotted on the same ordination space. For guilds, the designation using the different arrow line styles along the copiotrophy index is added independently using the results presented in Fig. 4. (A) NMDS results for the empirical communities, with color corresponding to the optimum of three clusters. (B) NMDS results for the modeled communities, with color corresponding to the optimum of three clusters. (C) NMDS results for the empirical communities, with color corresponding to five biogeographic regions, defined by latitude. (D) NMDS results for the modeled communities, with color corresponding to five biogeographic regions, defined by latitude. Because the sign of the axes is not meaningful, when comparing C and D, one could reverse the y-axis in one or the other in order to orient the subpolar vs. oligotrophic (and other) locations in the same order.

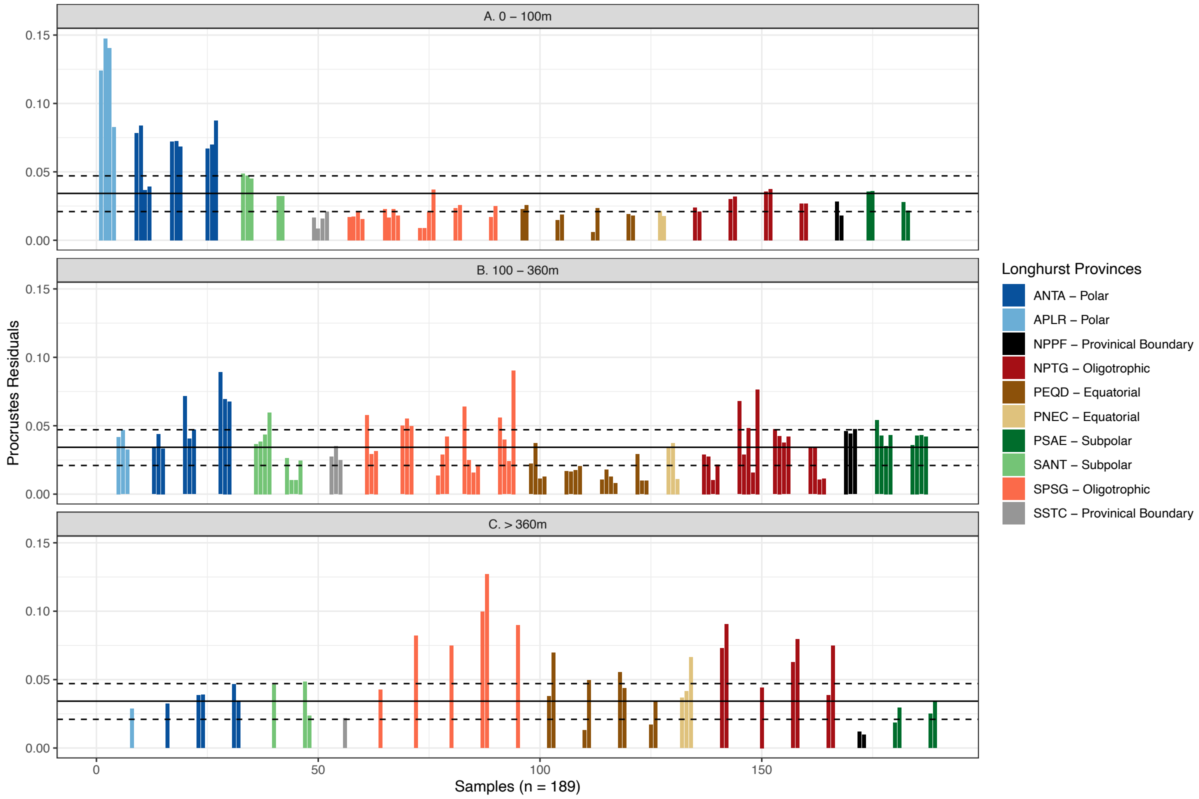


**Figure S17:** Residuals of the symmetric Procrustes rotation, highlighting the pointwise differences between the model and metabarcoding NMDS ordination, to aid in the interpretation of the regional and depth-structured misalignment between the model and molecular datasets. Each bar corresponds to a sample ($n$ = 189), with larger residuals indicating a greater misalignment between the two configurations and smaller residuals indicating a closer match. The solid line represents the median residual, and dashed lines highlight the lower (25%) and upper (75%) quantiles. The three panels represent the three depth bins: (A) 0 – 100 m, (B) 100 – 360 m, and (C) $>$ 360 m. The colors of each bar represent the corresponding Longhurst provinces of the sample.

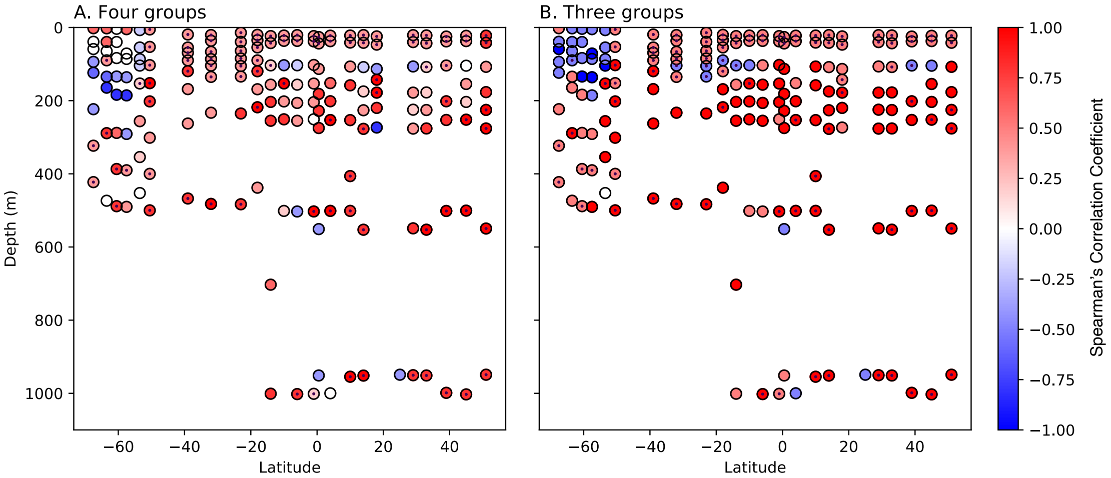


**Figure S18:** Rank correlation analysis results along the P16 transect. (A) Analysis for four comparable groups (see Materials and Methods and Fig. S14 for group descriptions). (B) Analysis for three comparable groups, combining two of the four in A. The color of the dot indicates the resulting Spearman's correlation coefficient. A small black dot in the center of the circle indicates that the model matched the observed dominant group (i.e., that the same group was ranked as first in both model and data). *Overall statistics:* Resulting means and standard deviations of the correlation coefficients were $0.40 \pm 0.42$ and $0.53 \pm 0.55$ for analysis of four and three groups, respectively, indicating overall positive correlation. The model matched the dominant type in 52% and 76% of the sample locations for four and three groups, respectively. The model most strongly departs from the data (exhibiting a strong negative correlation) in the Southern Ocean surface, where we compared data collected in the productive Austral summer to modeled annual averages. When neglecting this region (i.e., ignoring the sampling locations that are both below -50$^o$ latitude and above 200 m depth), the mean Spearman correlation coefficient was $0.67 \pm 0.40$ for the three groups, with the model matching the dominant group at 86% of the remaining locations.

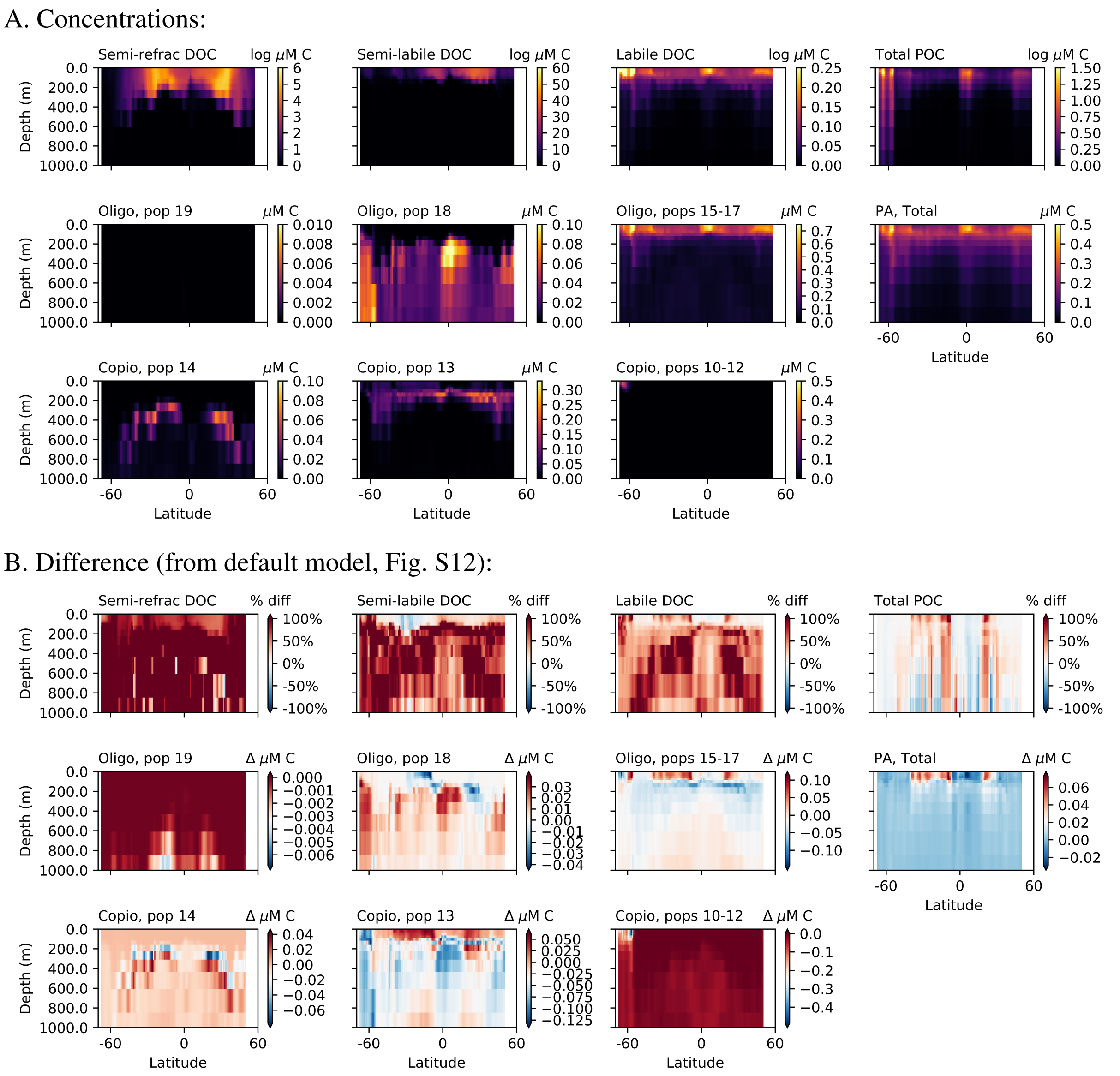


**Figure S19:** Results of a model experiment (showing just the Pacific transect) in which all heteroprokaryotic functional types (oligotrophs and copiotrophs) are consumed by one zooplankton grazer (which does not consume any phytoplankton). Though the magnitudes of DOC and biomass in this simulation vary from the default, the test is to see whether the mechanism of surface exclusion and the broad biogeographical patterns are still present, and this is indeed the case here. Thus, this experiment shows that the model results hold even without including the cell-size-based aspect of the grazing scheme.

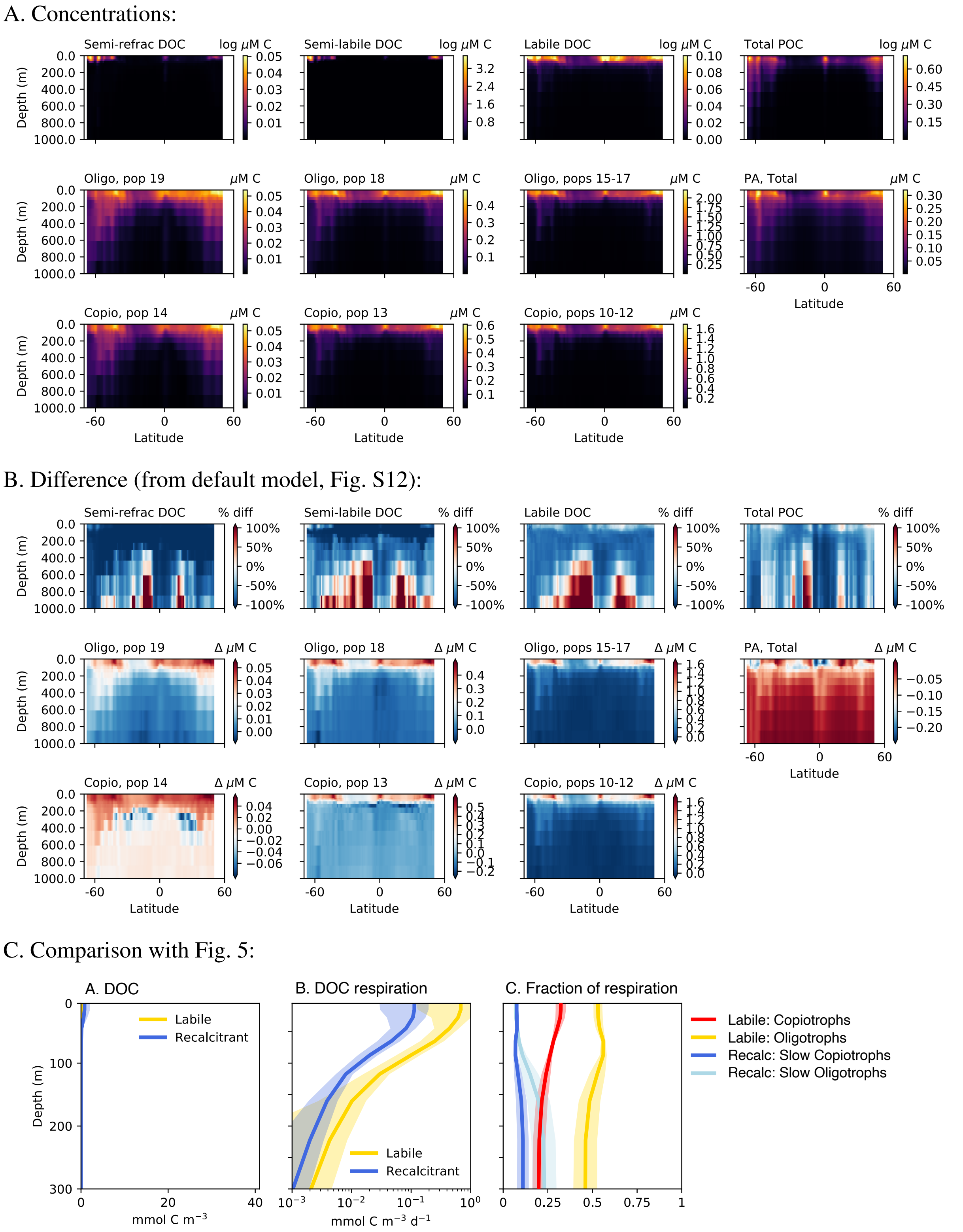


**Figure S20:** Results of a model experiment (showing just the Pacific transect) in which there is no explicit grazing of heteroprokaryotic by zooplankton. Instead, the quadratic mortality term for all functional type populations that were previously grazed (including all phytoplankton and DOM-consuming heteroprokaryotes) is increased 10-fold to account for grazing implicitly (*96*). This changes the ecosystem structure dramatically: it does not result in the surface exclusion of the slow-growing heteroprokaryotes consuming RDOM (semi-refractory and semi-labile DOC). Therefore, this experiment confirms that apparent competition (shared grazing) is the mechanism for the surface exclusion in the model.

A. Concentrations:

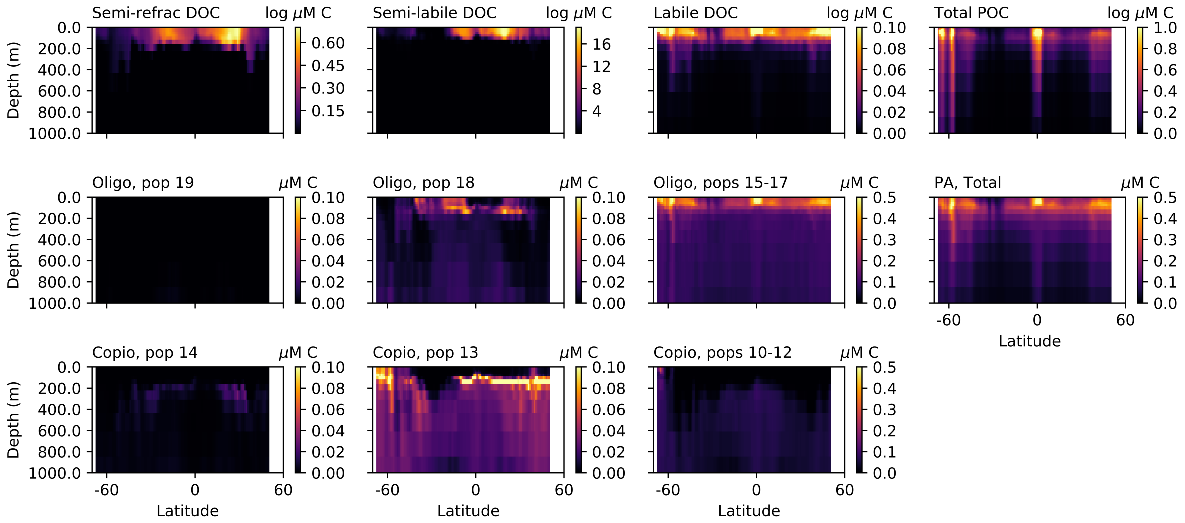


B. Difference (from default model, Fig. S12):

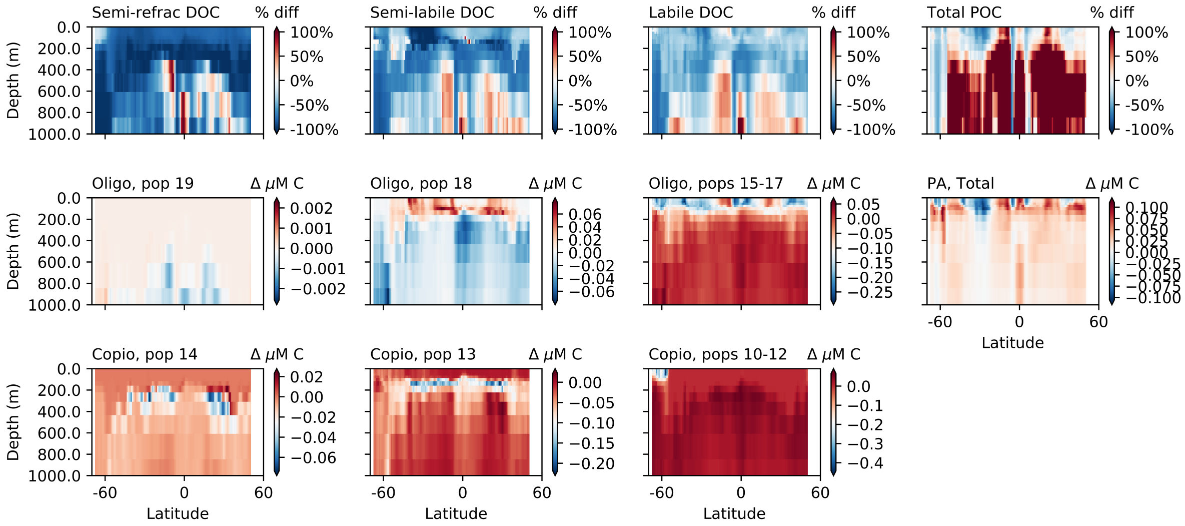


**Figure S21:** Results of a model experiment (showing just the Pacific transect) that serves to provide a baseline for a comparison of the next experiment in Fig. S22. Here, there is no excess hydrolysis of POM as a source of DOM ($\alpha = 1$), which means that total DOM production is less. In Fig. S22, we examine results with equal production of the two recalcitrant pools (semi-labile and semi-refractory pools), rather than following the lognormal distribution. We conducted this experiment because it is computationally easier to assure equal distribution to the two pools when there is not an additional source from POM. Because of the significantly lower supply of DOC, semi-labile DOC is depleted in some surface areas, and some of the consumers of RDOC (specifically, the oligotrophs) are able to survive at the surface in some locations).

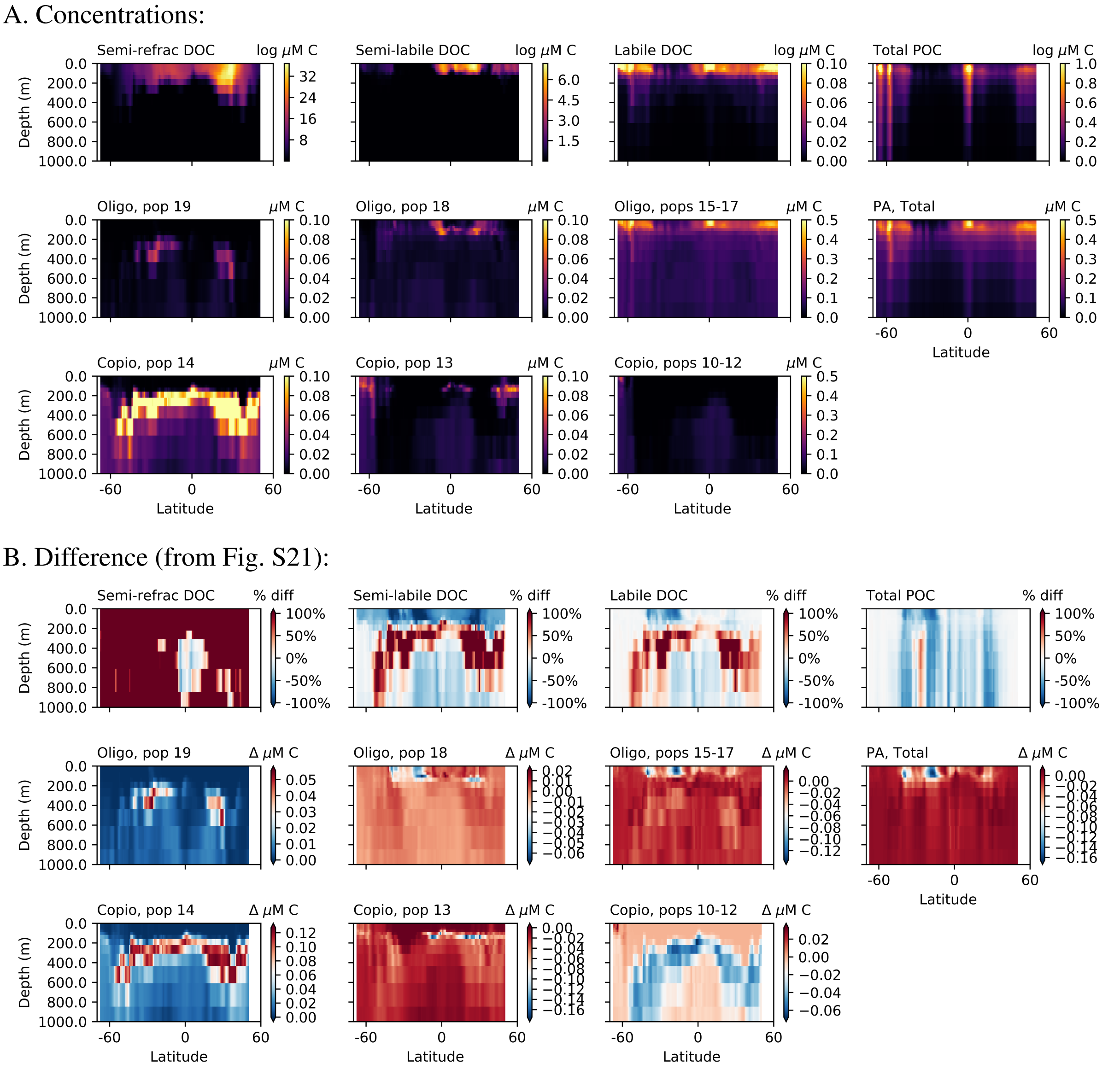


**Figure S22:** Results of a model experiment (showing just the Pacific transect) conducted to determine why the model predicts the greater ubiquity of the SAR324 analog relative to the SAR202 analog. In this experiment, the supply rate of the semi-labile and semi-refractory DOM is equal, rather than following the lognormal distribution, so that the only difference between the consumers is the difference in lability that sets the baseline maximum growth rate. The resulting biomasses of the semi-refractory consumers become much more significant, reflecting the higher substrate supply, but the biogeographical pattern remains. Therefore, the distinct biogeographical patterns between the semi-labile and semi-refractory consumers reflect differences in growth rate (i.e. lability), moreso than in substrate supply. The slower consumers of the semi-refractory DOC have biogeography that is more restricted to the center of the gyres compared with the more latitudinally widespread biogeography of the consumers of semi-labile DOC. From this, we hypothesize that SAR324 may grow more quickly than SAR202. Note that these results are directly compared with Fig. S21, where total DOC supply is much lower because $\alpha = 1$. This is because, to achieve the equal supply rates, we had to also turn off the excess hydrolysis of POM as a source of DOM ($\alpha = 1$). Note the different colorbar ranges for semi-refractory and semi-labile DOC here versus in Fig. S21.

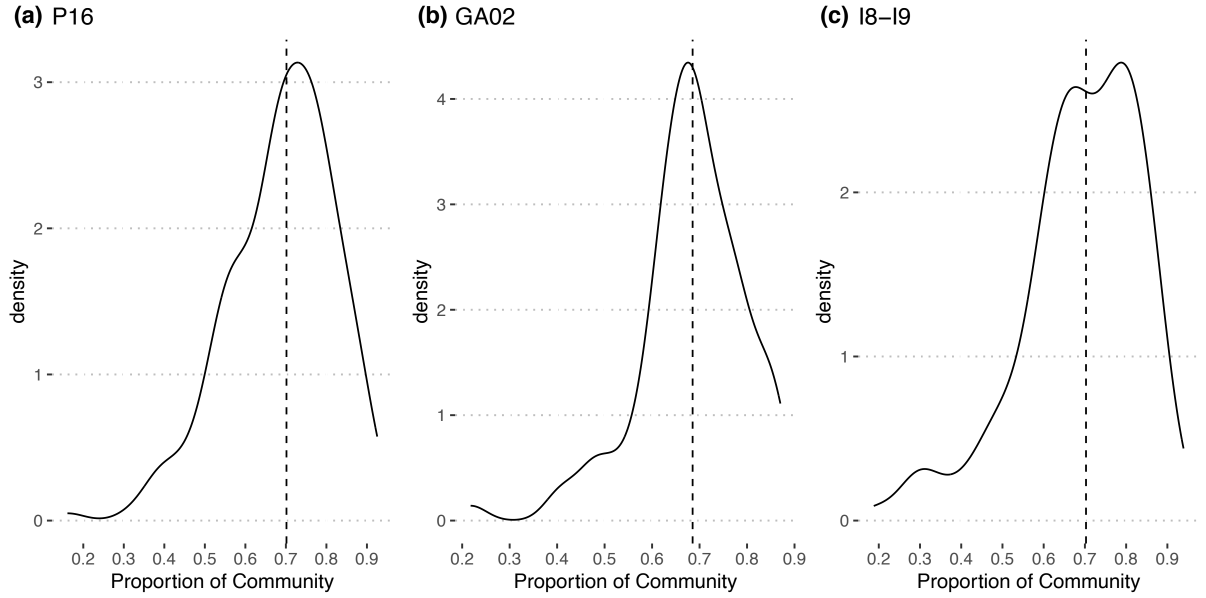


**Figure S23:** Proportion of the non-photosynthetic prokaryotic community covered by the genome matches to ASVs across the three transects, accounting for the relative abundance of the ASVs. For all three transects, most of the samples have roughly 70% coverage (median).

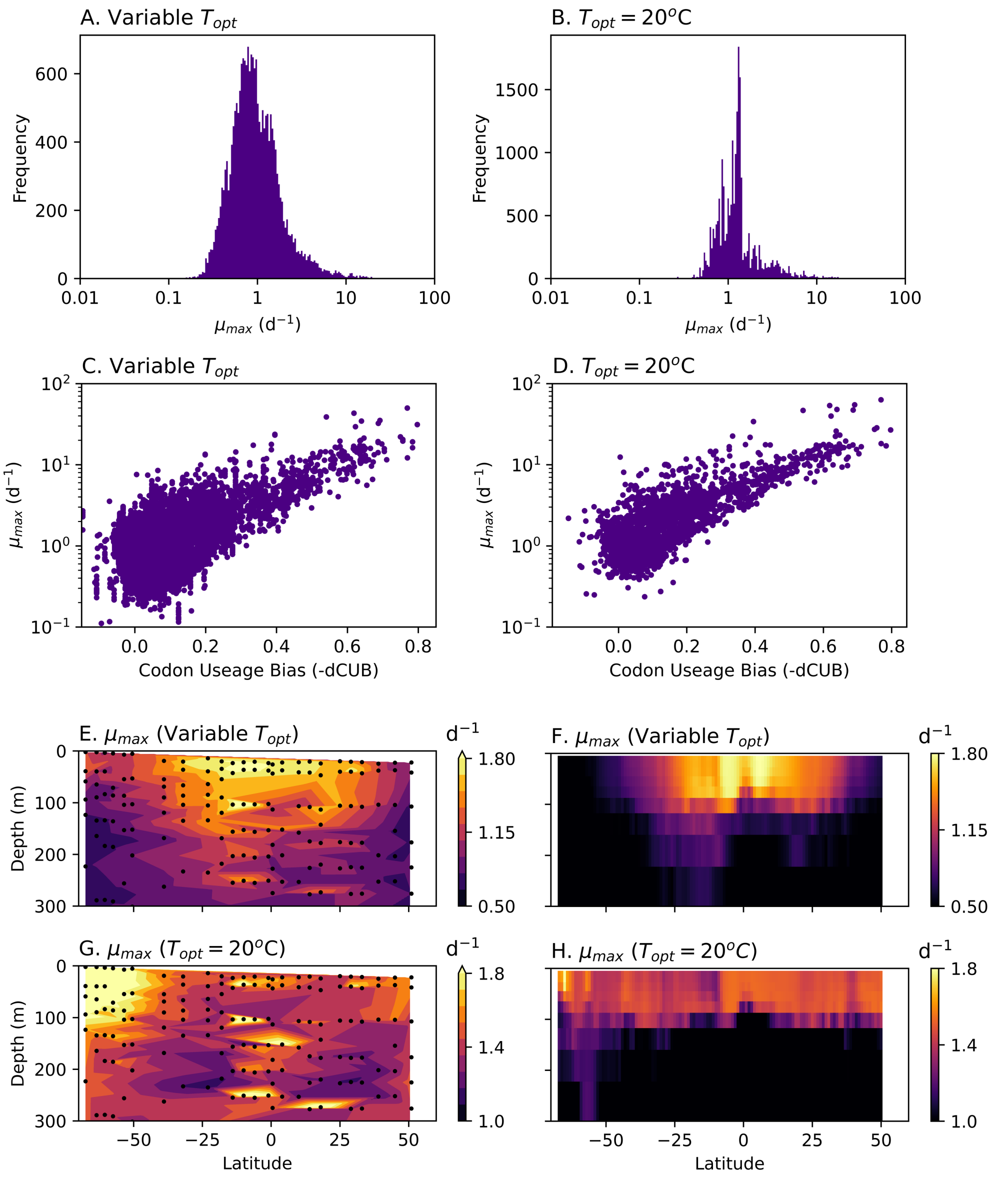


**Figure S24: Maximum growth rate estimates along the Pacific (P16) transect.** Maximum growth rate ($\mu_{max}$) is estimated using codon useage statistics (*19*). (A) Distribution of $\mu_{max}$ for each ASV when the optimum temperature is set by the location in which that ASV exhibits the top 1% of its abundance. (B) Distribution of $\mu_{max}$ when the optimum temperature for each ASV is set to 20$^o$C. (C) and (D) Relationship between the predominant codon useage bias statistic (dCUB) and maximum growth rate ($\mu_{max}$) estimates. More bias (lower values of dCUB) results in a higher maximum growth rate estimate, but there is additional scatter, indicating that other factors contribute to the maximum growth rate in the algorithm. $\mu_{max}$ is estimated using dCUB as well as other codon useage statistics (*19*). (E–H) Community-averaged (relative-abundance weighted) heteroprokaryotic maximum growth rates (2D version) along the Pacific (P16) transect in data and model (DOM-consuming community only). (E) and (F) $\mu_{max}$ when the optimum temperature for each ASV is set by the location in which that ASV exhibits the top 1% of its abundance for the data, and when the water temperature modifies $\mu_{max}$ in the model following Eqn. S5. (G) and (H) $\mu_{max}$ when the optimum temperature for each ASV and the temperature that modifies $\mu_{max}$ in the model is set constant at 20$^o$C. For the model, we here look just at the average of the DOM-consuming community in order to be able to illustrate the transition from oligotrophy to copiotrophy at the surface in the T = 20$^o$C. The smaller latitudinal surface gradient in the model demonstrates that the modeled $\Delta\mu_{max}$ between oligotrophs and copiotrophs is smaller than the empirical estimates, although any magnitude of difference (i.e. tradeoff between $\mu_{max}$ and affinity) allows the model to reproduce the transition of the functional types.

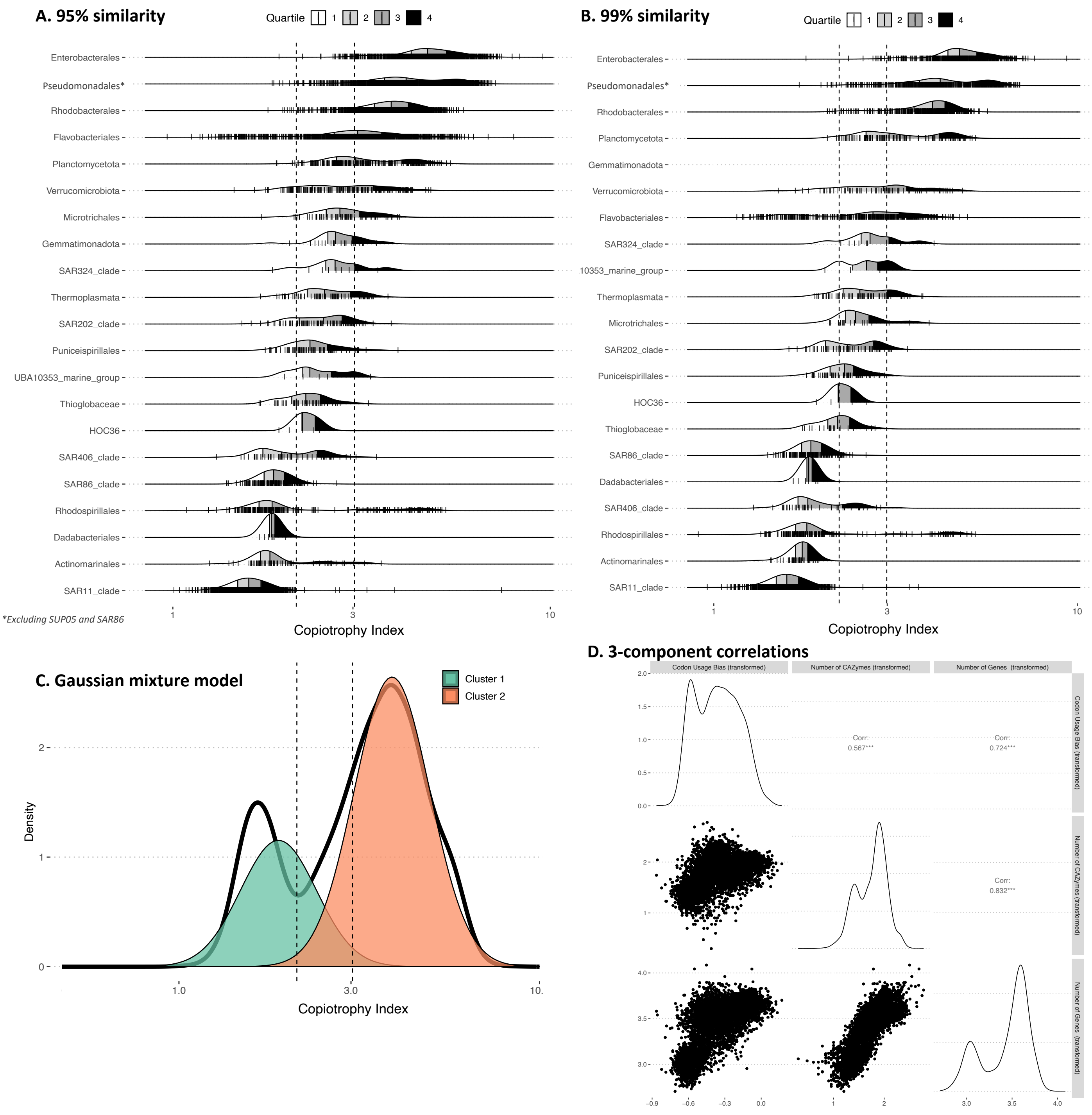


**Figure S25:** The genome-based copiotrophy index results and analysis. Results for (A) 95% similarity, (B) 99% similarity, (C) Gaussian mixture model showing the emergence of two lognormal distributions, and (D) correlations between each of the three components of the copiotrophy index: the transformed codon useage bias, the number of carbohydrate-active enzymes (CAZymes), and the number of genes.

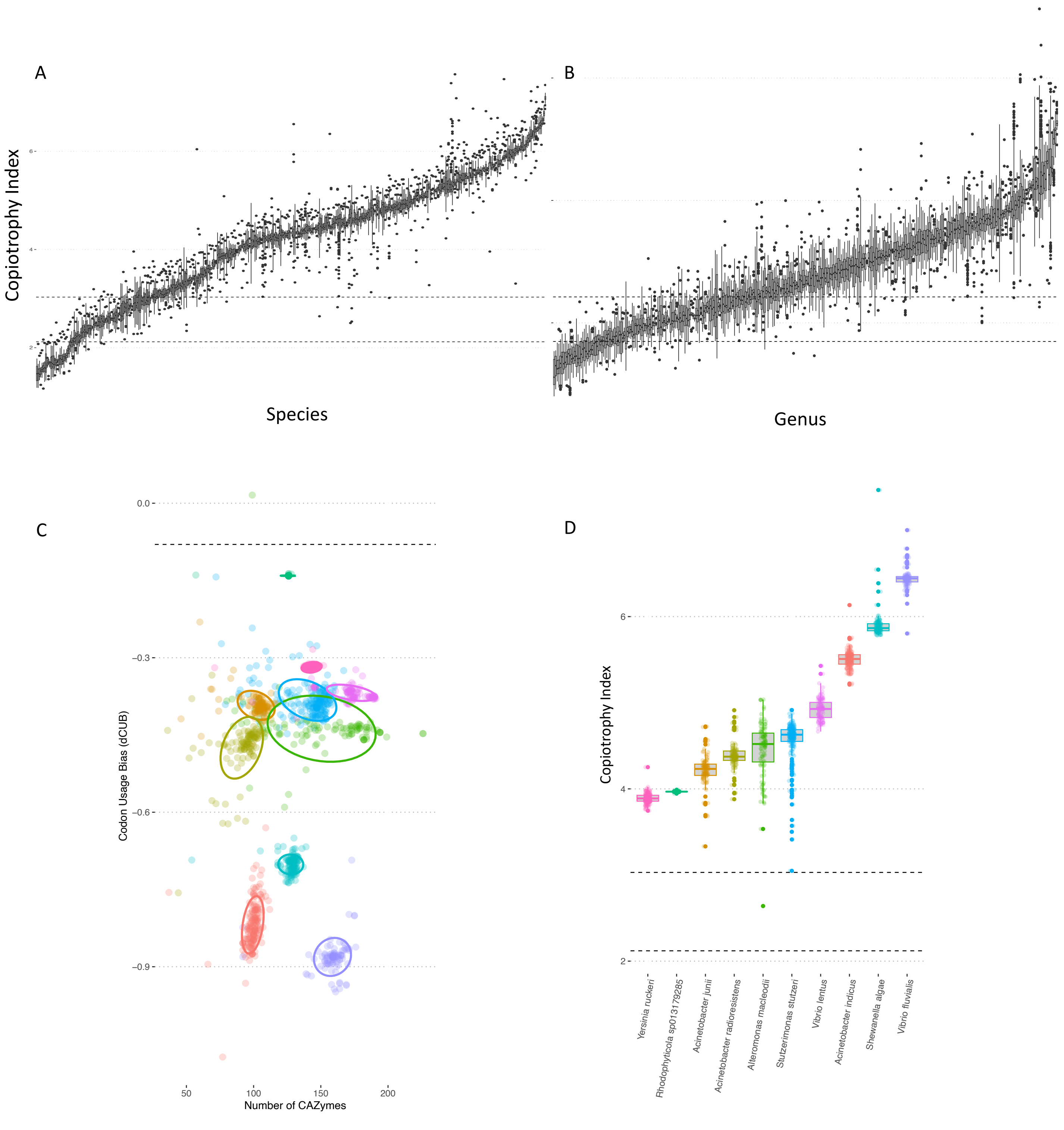


**Figure S26:** Analysis showing consistency of the copiotrophy index calculated for: (A) within-species strains and (B) within-genus strains. The within-species analysis includes all species with at least 10 representatives in GTDBv220 that also have a least one genome match and a taxonomic annotation. (C and D) Further analysis of within-species patterns, using a subset of the species with hits on our transect that have at least 100 (rather than 10) genomes in GTDBv220. Panel C shows the relationship between dCUB and CAZy, two of the three factors for the copiotrophy index. Panel D shows that the resulting copiotrophy index range within each species is generally quite narrow, even when the dCUB or nCAZy range is wider.

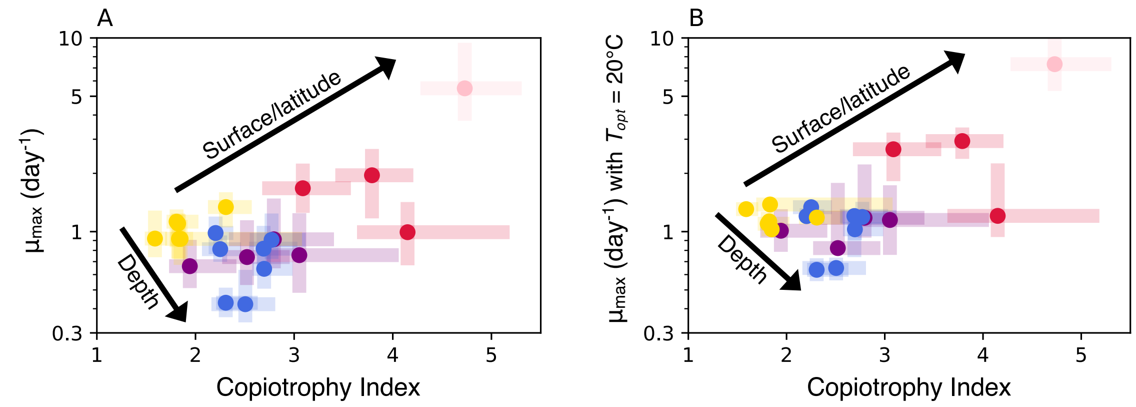


**Figure S27: The relationship between the copiotrophy index and the maximum growth rate ($\mu_{max}$) for each guild.** The median values and interquartile ranges are plotted for each guild. (A) Relationship when $\mu_{max}$ reflects the optimum temperature for each ASV, set by the location in which that ASV exhibits the top 1% of its abundance (B) Relationship when the optimum temperature for each ASV is set to 20$^o$C. When controlling for temperature (panel B), the differences in maximum growth rates between the low-latitude surface guilds (yellow dots) and the deep guilds (blue dots) are less, though two of the deep guilds still exhibit the lowest medians.

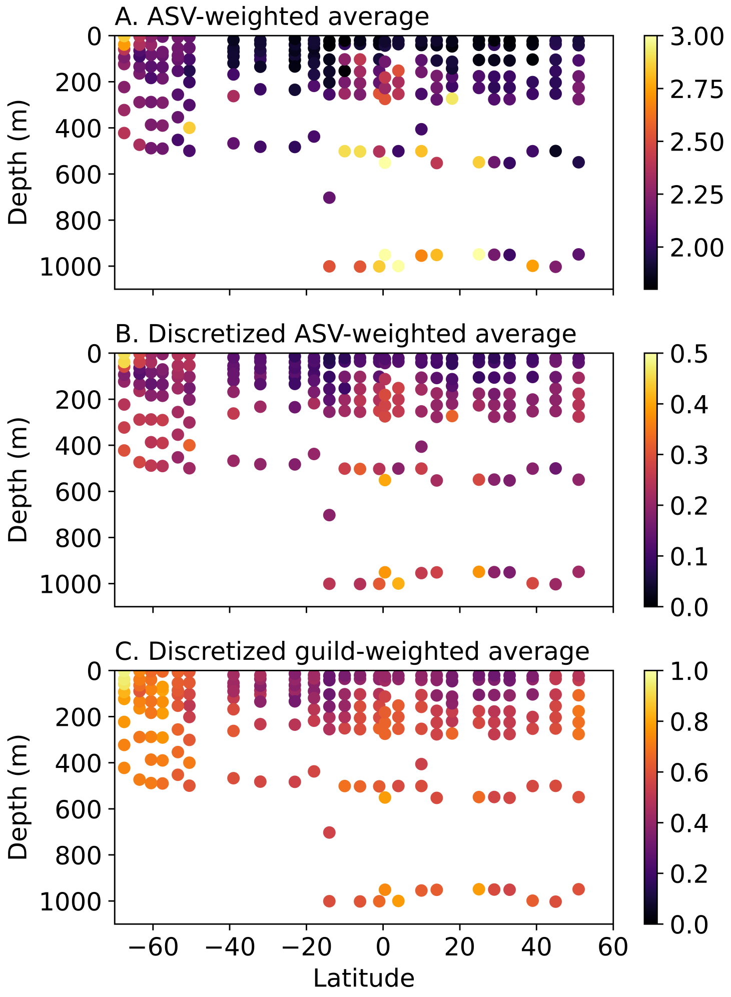


**Figure S28:** The copiotrophy index at the community level along the P16 transect. (A) The relative-abundance weighted average of the copiotrophy index for the individual ASVs for all non-photosynthetic prokaryotes. (B) As in panel A, but with the copiotrophy index discretized in order to compare most directly with the global model by assigning a value of 0 to oligotrophs and 1 to slow copiotrophs and copiotrophs (matching the model configuration) according to the copiotrophy index of each ASV. (C) As in panel B, but at the guild level, according to the median copiotrophy index of the guild. Panel C is illustrated in the main text in an interpolated format (Fig. 4A). All show a similar pattern, with the high-latitude copiotrophs more copiotrophic than the "slow copiotrophs," consistent with the ranking in Fig. 3.

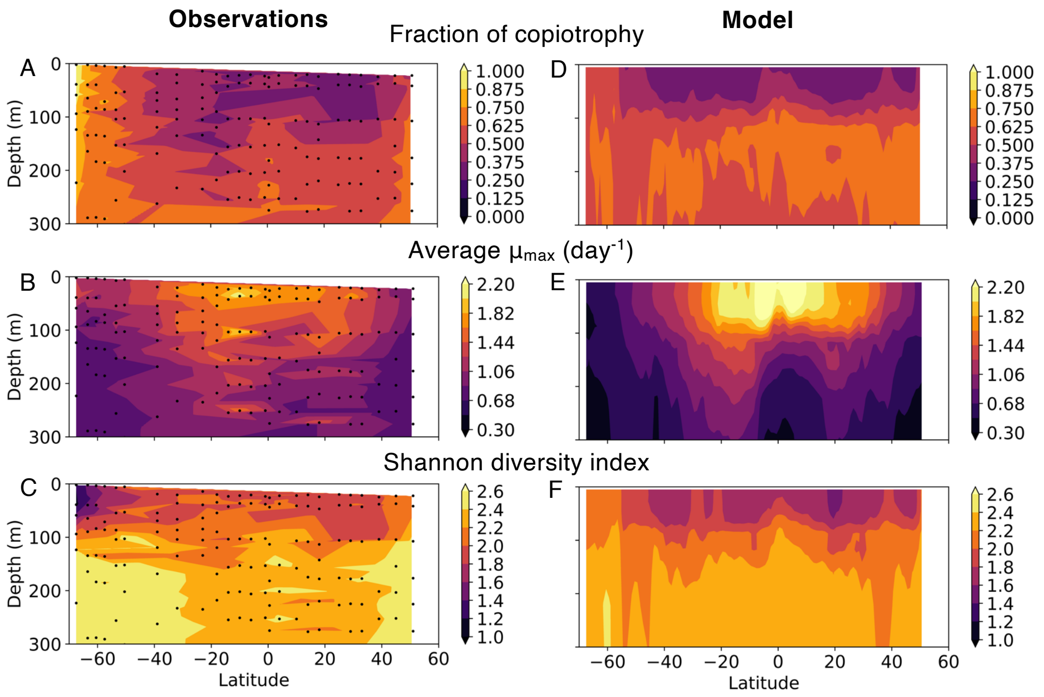


**Figure S29: Synthesis of observed and modeled heteroprokaryotic community patterns along the P16 transect.** (A and D) Fraction of the heteroprokaryotic community classified as copiotrophic. (B and E) Community-averaged maximum growth rate ($\mu_{max}$) estimate (*19*). (C and F) Shannon diversity index, calculated with respect to the taxonomic guilds (C) and the modeled functional types (F) to allow for an 'apples-to-apples' comparison.

**Table S1:** Guild name (short version), keywords for ASV search, the number of ASVs in each guild (for the P16 transect), the number of ASV-matched genomes for each guild (for the P16 transect), the resulting copiotrophy index (CI) median and interquartile range (IQR) values, resulting genome-based maximum growth rate ($\mu_{\max}$) estimates using the codon-useage-bias-based gRodon2 algorithm (*19*). The copiotrophy index (CI) median values differentiating the clusters (the vertical dashed black lines in Fig. 4 in the main text) are 2.12 and 3.03.

| Guild Name | grep search keywords | # of ASVs | # of genomes | CI median | CI IQR | gRodon2 $\mu_{\max}$ median | gRodon2 $\mu_{\max}$ IQR |
|---|---|---|---|---|---|---|---|
| Enterobacterales | o__Enterobacterales | 481 | 978 | 4.73 | 1.03 | 5.49 | 5.70 |
| Pseudomonadales* | Alcanivoracaceae, Cellvibrionaceae, Endozoicomonadaceae, Halieaceae, Halomonadaceae, KI89A_clade, Litoricolaceae, Marinobacteraceae, Moraxellaceae, Nitrincolaceae, Oceanospirillaceae, Oleiphilaceae, OM182_clade, Porticoccaceae, Pseudohongiellaceae, Pseudomonadaceae, Saccharospirillaceae, Spongiibacteraceae, g__Alkalimarinus | 1375 | 1243 | 4.15 | 1.55 | 0.99 | 0.75 |
| Rhodobacterales | o__Rhodobacterales | 477 | 1191 | 3.79 | 0.79 | 1.95 | 1.50 |
| Flavobacteriales | o__Flavobacteriales | 2453 | 1220 | 3.09 | 0.90 | 1.67 | 1.00 |
| Planctomycetota | p__Planctomycetota | 1656 | 230 | 3.05 | 1.35 | 0.76 | 0.76 |
| Verrucomicrobiota | p__Verrucomicrobiota | 1143 | 152 | 2.79 | 1.07 | 0.91 | 0.84 |
| Microtrichales | o__Microtrichales | 177 | 66 | 2.77 | 0.57 | 0.91 | 0.52 |
| Gemmatimonadota | p__Gemmatimonadota | 224 | 24 | 2.70 | 0.41 | 0.64 | 0.35 |
| SAR324_clade | p__SAR324_clade | 545 | 9 | 2.69 | 0.49 | 0.82 | 0.40 |
| Thermoplasmata | c__Thermoplasmata | 533 | 77 | 2.52 | 0.68 | 0.74 | 0.61 |
| SAR202_clade | o__SAR202_clade | 1626 | 91 | 2.50 | 0.65 | 0.42 | 0.29 |
| Puniceispirillales | o__Puniceispirillales | 531 | 70 | 2.31 | 0.44 | 1.34 | 0.55 |
| UBA10353_marine_group | o__UBA10353_marine_group | 345 | 5 | 2.31 | 0.42 | 0.43 | 0.15 |
| Thioglobaceae | f__Thioglobaceae | 105 | 48 | 2.25 | 0.40 | 0.81 | 0.26 |
| HOC36 | o__HOC36 | 161 | 5 | 2.20 | 0.19 | 0.98 | 0.43 |
| SAR406_clade | p__Marinimicrobia_(SAR406_clade) | 4111 | 54 | 1.94 | 0.68 | 0.66 | 0.40 |
| SAR86_clade | f__SAR86_clade | 617 | 156 | 1.85 | 0.23 | 0.92 | 0.48 |
| Rhodospirillales | o__Rhodospirillales | 2446 | 201 | 1.83 | 1.39 | 0.92 | 0.51 |
| Dadabacteriales | o__Dadabacteriales | 18 | 7 | 1.82 | 0.06 | 1.10 | 0.52 |
| Actinomarinales | o__Actinomarinales | 95 | 57 | 1.81 | 0.32 | 1.13 | 0.47 |
| SAR11_clade | o__SAR11_clade | 3202 | 981 | 1.59 | 0.23 | 0.92 | 0.55 |

*Excluding SUP05 and SAR86 (hence the long list of keywords).

**Table S2:** Parameter values for the heteroprokaryotic functional types and DOM lability classes. The distribution weight follows a lognormal distribution and governs the relative production of each DOM class from total DOM production in the model. Maximum growth rate ($\mu_{max}$) values are for the reference temperature of 20$^{o}$C. Together, $\mu_{max}$ and the half-saturation coefficient ($k_{DOC}$) scale allometrically and govern the uptake affinity for the carbon component of each DOM class by each functional type. The half-saturation coefficients for the other organic elemental pools follow a constant stoichiometry for marine bacterial biomass (*80*).

| DOM class | Weight | Copio $\mu_{max}$ (d$^{-1}$) | Oligo $\mu_{max}$ (d$^{-1}$) | Copio $k_{DOC}$ ($\mu$M C) | Oligo $k_{DOC}$ ($\mu$M C) |
|---|---|---|---|---|---|
| DOM 1 (most labile) | 0.0018 | 32 | 22 | 7.7 | 3.5 |
| DOM 2* | 0.15 | 5.6 | 4.0 | 1.4 | 0.62 |
| DOM 3* | 0.69 | 1.0 | 0.71 | 0.24 | 0.11 |
| DOM 4* | 0.15 | 0.18 | 0.13 | 0.043 | 0.020 |
| DOM 5 (least labile) | 0.0018 | 0.029 | 0.022 | 0.0077 | 0.0035 |

*The three POM consuming functional types match the values of the copiotrophs for the three middle DOM lability classes.
